# scTail: precise polyadenylation site detection and its alternative usage analysis from reads 1 preserved 3’ scRNA-seq data

**DOI:** 10.1101/2024.07.05.602174

**Authors:** Ruiyan Hou, Yuanhua Huang

## Abstract

Three-prime single-cell RNA-seq (scRNA-seq) has been widely employed to dissect the variability of cellular transcriptomes, while only the cDNAs on reads 2 are routinely used, including to analyze polyadenylation sites (PAS). However, despite of high sequencing noise, we found the cDNAs on reads 1 are highly informative in precisely detecting PAS. Here, we further develop a computational method, scTail, to identify PAS using reads 1 and quantify its expression leveraging reads 2, which enables effective detection of alternative PAS usage (PAU). When compared with other methods, scTail detects PAS more sensitively and precisely. With various experimental data sets, we demonstrated that the combination usage of scTail and BRIE2 can discover differential alternative PAS usage in various biological processes including cell types in human intestinal, disease status of esophageal squamous cell carcinoma, and time point of mouse forelimb histogenesis, revealing critical insights in PAS regulations.

## Introduction

With the assistance of RNA polymerase II, DNA is transcribed into pre-mRNA, which subsequently undergoes processing to become mature mRNA. The processing stages involve adding a cap, splicing, as well as cleavage and polyadenylation (CPA) [1]. CPA is essential for transcription termination and comprises the endonucleolytic cleavage of a nascent transcript and polyA tail synthesis. The site of cleavage at which polyA tail is added is called polyadenylation site (PAS). One gene can selectively utilize different PASs, termed alternative polyadenylation (APA), which generate RNA isoforms with different 3’ untranslated regions (3’ UTRs) or protein coding sequences [2]. APA plays essential roles in regulating mRNA stability, translation efficiency, and subcellular localization [3]. More studies evidenced the importance of PAS regulation for different physiological and pathological conditions, such as development [4, 5], oncogenesis [6, 7] and inflammation [8]. Furthermore, the 3’ UTR of therapeutic mRNAs may play a pivotal role in maintaining their intrinsic stability and modulating the intracellular dynamics [9], therefore understanding the PAS selection and 3’ UTR function in natural contexts may also provide translational insights.

The emergence of RNA sequencing (RNA-seq) techniques has facilitated the examination of the transcriptome. Although it was not designed for detecting PAS, several studies still utilized RNA-seq data to profile APA by checking reads coverage changes at PASs [10] or taking advantage of annotated PAS [11]. On the other hand, some experimental protocols have enabled to capture 3’ ends of mRNAs for direct identifying PAS, such as PolyA-seq [12], PAS-seq [13], 3’READS [14] and QuantSeq REV [15]. In recent years, single-cell RNA sequencing (scRNA-seq) has become a commonly used technology for analyzing gene expression at a cellular resolution. Protocols that utilize oligo(dT) priming for cDNA generation and library construction were called 3’ tag-based scRNA-seq, such as 10x Genomics’s chromium single cell 3’ solution (now a popular commercial choice) [16], Drop-seq [17] and CEL-seq [18]. Multiple bioinformatic methods were also developed to detect and analyze PAS based on the 3’ tag-based scRNA-seq, including scAPA [19], Sierra [20], scDaPars [21], MAAPER [22], scAPAtrap [23], SCAPTURE [24], SCAPE [25] and Infernape [26]. Interestingly, Infernape also demonstrated its capability in support of PAS analysis from spatial 3’ RNA-seq data.

However, all of these methods only use reads2 in single cell 3’ RNA-seq (e.g., from 10x Genomics), which are fragmented and cannot reach PAS precisely. According to our observation, the fragmentation does not affect the sequence close to poly(dT)VN that can be captured by the cDNA in reads1 (if sequenced and preserved), making it possible to detect polyadenylation site at a near single nucleotide resolution (Fig. 1A). As such information is conventionally overlooked, majority of 3’ scRNA-seq data sets from 10x Genomics platform do not contain cDNA sequences in reads 1 by either being trimmed or not sequenced. Nevertheless, there are still a good number of public 3’ 10x Genomics datasets on the GEO repository retaining such information, therefore providing an opportunity to study alternative polyadenylation usage coupled with gene expression at a single cell level without extra cost. These observations were well supported by a recent report by Fu and colleagues that cDNAs on reads 1 can be informative in PAS detection despite the sub-optimal base calling quality, presumably due to higher sequencing errors in homopolymer stretches [27]. However, with the relatively low sequencing quality, it is unclear how to balance the reads mapping quality and PAS detection sensitivity. Potentially, combining with the sequence modeling of the PAS, e.g., with deep neural networks, may help filter false PASs hence improving the specificity while maintaining a high sensitivity. Moreover, a coherent method suite for PAS detection, expression counting, and differential usage identification is still urgently demanded to popularize the PAS analysis with existing scRNA-seq solutions; ideally, it should seamlessly support common single-cell analysis ecosystems (e.g., Scanpy).

**Figure 1.**
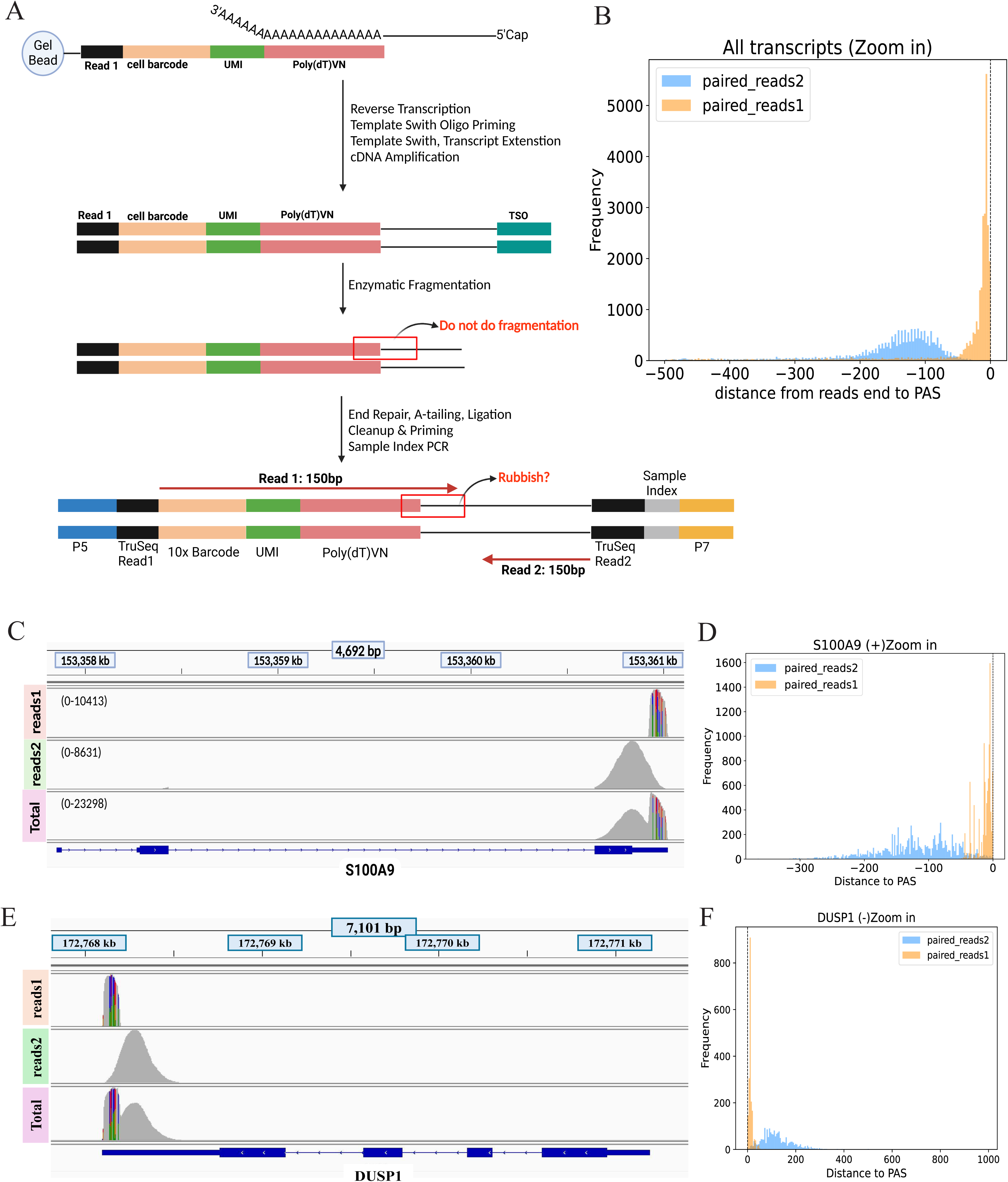
Evidences of extra cDNA from reads 1 is better than reads 2 to detect PAS. **(A)** A flow chart of the 3’ scRNA-seq gene expression library construction (10x Genomics). **(B)** Histogram displays the most frequent distance from the end of reads 1 and reads 2 to annotated PAS for each transcript. **(C)** Screenshot from IGV shows the coverage of reads 1, reads 2, and total reads for forward strand gene S100A9. **(D)** Histogram shows the distance between reads 1 or 2 and annotated PAS for forward strand gene S100A9. **(E)** Screenshot from IGV shows the coverage of reads 1, reads 2, and total reads for reverse strand gene DUSP1. **(F)** Histogram shows the distance between reads 1 or 2 and annotated PAS for reverse strand gene DUSP1.

In this work, to resolve the above challenges, we introduce an all-in-one stepwise method scTail to analyse alternative PAS usage from reads 1 preserved 3’ scRNA-seq data. Specifically, it leverages the cDNA in reads 1 to precisely detect PAS that benefited from including a deep learning-based classifier for noise filtering, whose expression level can be further estimated by probabilistically assigning reads 2. Its high accuracy is first evidenced by the consistency of known PAS sequence motifs and existing annotations. With the high sensitivity empowered by reads 1, we further demonstrated that PAS analyses, especially the alternative usage, can reveal remarkable biological insights, including in cell type characterization, intestinal differentiation, mouse forelimb development, esophageal squamous cell carcinoma, and the regulation of RNA binding proteins. Overall, these analyses well evidence the important roles that alternative PAS usage plays in diverse tissues and processes.

## Results

### Design of scTail for PAS analysis with both reads 1 and 2

To illustrate that the cDNA in reads 1 (termed reads 1 below for simplicity) is informative in detecting PASs, we examined an example dataset profiled by 10x Genomics (PBMC sample from the ESCC dataset, see below) by setting higher mapping error tolerance and anchoring with read 2. First, we calculate the distance between the UMI end and the annotated PAS in GENCODE. Overall, the histogram in Fig. 1B shows the distribution of the most frequent distance between the UMI ends to PAS for all 252,835 annotated transcripts. It illustrates that read 1’s ends are tightly close to the annotated PAS while read 2’s ends are around 12 times further (median: -11 vs -132) and the former has a sharper distribution; interestingly, the distance between reads 1 and reads 2 is approximately to the length of fragmentation (Supplementary Fig. 1A). Here, we selected two example genes (S100A9 and DUSP1) that only have one annotated transcript. As screenshots from the integrative genomics viewer (IGV) shown in Fig. 1C and Fig. 1E, most of reads 1 are more closely located to the annotated PAS with a narrower peak compared to that of reads 2, which is more evident if only looking at the read ends instead of the base coverage (Fig. 1D, F, Supplementary Fig. 1B, C). Of note, reads 1 indeed present more mismatches as shown in the IGV plot (Fig. 1C), but are still highly informative in detecting PAS if we set a higher error tolerance.

To support and popularize the PAS analysis from the existing 3’ scRNA-seq solutions (10x Genomics) with reads 1, we developed an all-in-one stepwise computational method called scTail. In brief, scTail takes an aligned bam file from STARsolo (with higher tolerance of low-quality mapping; see Methods) as input and returns the detected PASs and a PAS-by-cell expression matrix. It mainly comprises two major steps: 1) call peaks by leveraging paired reads 1 (peaks refer to PAS; Fig. 2A). 2) quantify expression for each peak called at the first step at single-cell resolution (Fig. 2B). As shown in Fig. 2A, aligned bam file contains four kinds of reads: paired reads 1, paired reads 2, unique reads 1 and unique reads 2. First, the terminate positions of paired reads 1 were selected to identify the putative PAS via clustering with paraclu [28], which also has a module to remove clusters according to the technical thresholds (minimum cluster UMI count: 50 by default; maximum cluster length: 100 bp by default; minimum density increase: 0 by default, via paraclu-cut.sh). In addition, a deep learning neural network was embedded in scTail to further filter false positive artifacts (see next paragraph). Second, due to the limitation of the low mapping rate for reads 1, we took advantage of total reads 2 (i.e., both paired and unique reads 2) to quantify the PAS expression by assigning reads 2 to the detected PASs via modeling the distribution of the fragment size. Specifically, this fragment size distribution is generally assumed the same within a dataset but varies among datasets, and it can be empirically estimated from the distances of 3’ ends between paired reads 1 and 2 by fitting a log-normal distribution, ideally focusing on genes with one single PAS. This fragment size distribution also represents the distance of reads 2 ends and their true PAS (nearly the reads 1 ends), hence can be used to calculate the likelihood of a read 2 being assigned to a certain PAS (3’ end of the detected PAS cluster). Then, within each gene, we can assign reads 2 to the PAS that has the highest likelihood (if there are multiple) and passes a threshold (see Methods). Finally, a user-friendly and scalable output file format will present UMI counts for each PAS in each cell (Fig. 2B).

**Figure 2.**
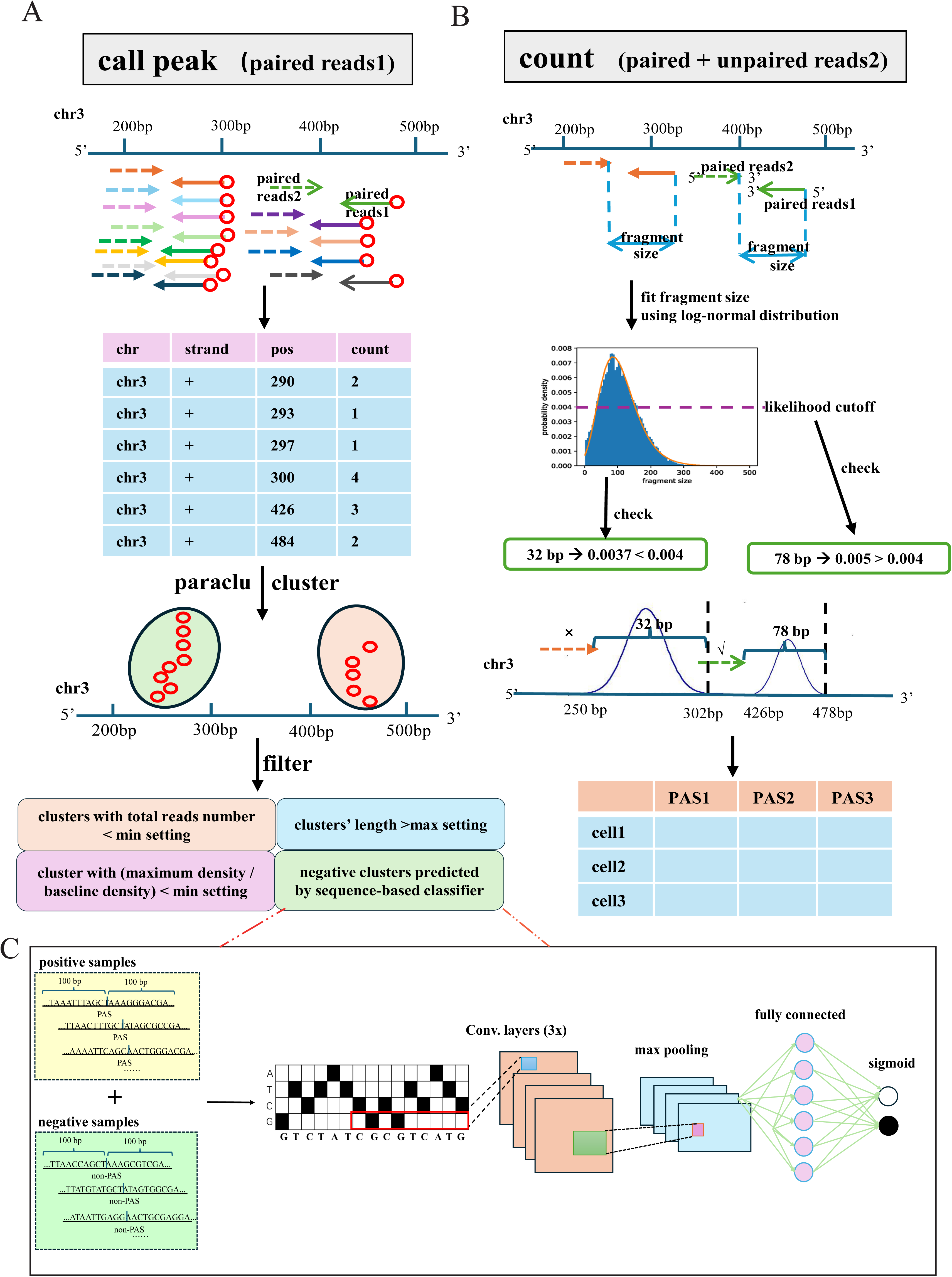
Developing scTail to identify PAS from 3’ tag-based scRNA-seq. **(A)** Outline of the first step of scTail. The purpose of this step is to get the position of PAS from aligned bam file. In the real situation, calling peak was going on for each sample (one bam file). Then peaks of all samples were merged. The red circle is the terminate position of reads1. **(B)** Schematic of the second step of scTail. An aligned bam file was inputted and a sparse matrix (h5ad file) was obtained as output. **(C)** Flowchart of deep neural network embedded into scTail to filter low-quality putative PASs from sequence.

In scTail, the classifier plays an essential role in the filtering step to rule out the false positive peaks, for example produced by internal priming of oligo(dT) at A-rich sequences or aberrant read augment during library construction [24] (Fig. 2C). Here, we employed a convolutional neural network (CNN; architecture detail in Methods) as the classifier considering the strong sequence patterns around PAS [29]. To train the model, positive PASs were collected from GENCODE and three other databases: PolyA DB3, PolyA-seq, and PolyASite, while negative PASs were randomly picked from either the intergenic region that did not overlap with positive PASs or transcriptional start sites (TSSs). The sequence between upstream 100 bp and downstream 100 bp of sample position was extracted and encoded with a one-hot coding method as the input features to the CNN model. Finally, our trained model can predict the putative PAS produced by paraclu in a new dataset as a positive or negative PAS, where only the positively predicted ones are kept for downstream analysis.

### scTail identifies PAS accurately and sensitively

As mentioned above, scTail embedded a pre-trained sequence model to remove the false positive clusters, which enabled us to further evaluate the reliability of the detection by examining the supervised performance metrics and learned sequence motifs. For the classification performance in cross-validation, the sequence model achieves high performance in both sensitivity and specificity, with the area under the receiver operating characteristic curve (AUROC) of 0.985 and 0.979 in human and mouse, respectively (Fig. 3A). To ensure the model captures the interpretable features, the maximum activation seqlet [30] in the human model was utilized to visualize individual convolutional filters at the second layer by leveraging the K562 dataset as a test dataset. Some of the most frequently observed patterns were the AAUAAA motif located 25-35 bp upstream of the cleavage site, as well as downstream GU-rich, U-rich, and G-rich motifs. All of these motifs and their activated positions are highly consistent with previous reports [2], indicating that our sequence model can identify reliable PASs (Fig. 3B, Supplementary Fig. 2). After filtering the potential false positives, Fig. 3C shows the majority of PASs (73.6%) detected from the K562 cell line are annotated whereas the other PASs (26.2%) identified by scTail are novel. As expected, most PASs (8,810 out of 13,937) were mapped to 3’ UTR, while a substantial portion of PASs was also mapped to exon (1,525) and intron (3,540), presumably as a result of alternative usage of polyadenylation site (Fig. 3D). Next, we plotted the nucleotides profile around PASs to assess the precision of detected PASs and observed canonical nucleotide distributions in total, annotated and unannotated PASs identified by scTail, exhibiting consistency with known PAS motifs (Fig. 3E, Supplementary Fig. 3), hence consolidating the accuracy of scTail in PAS detection including novel PAS.

**Figure 3.**
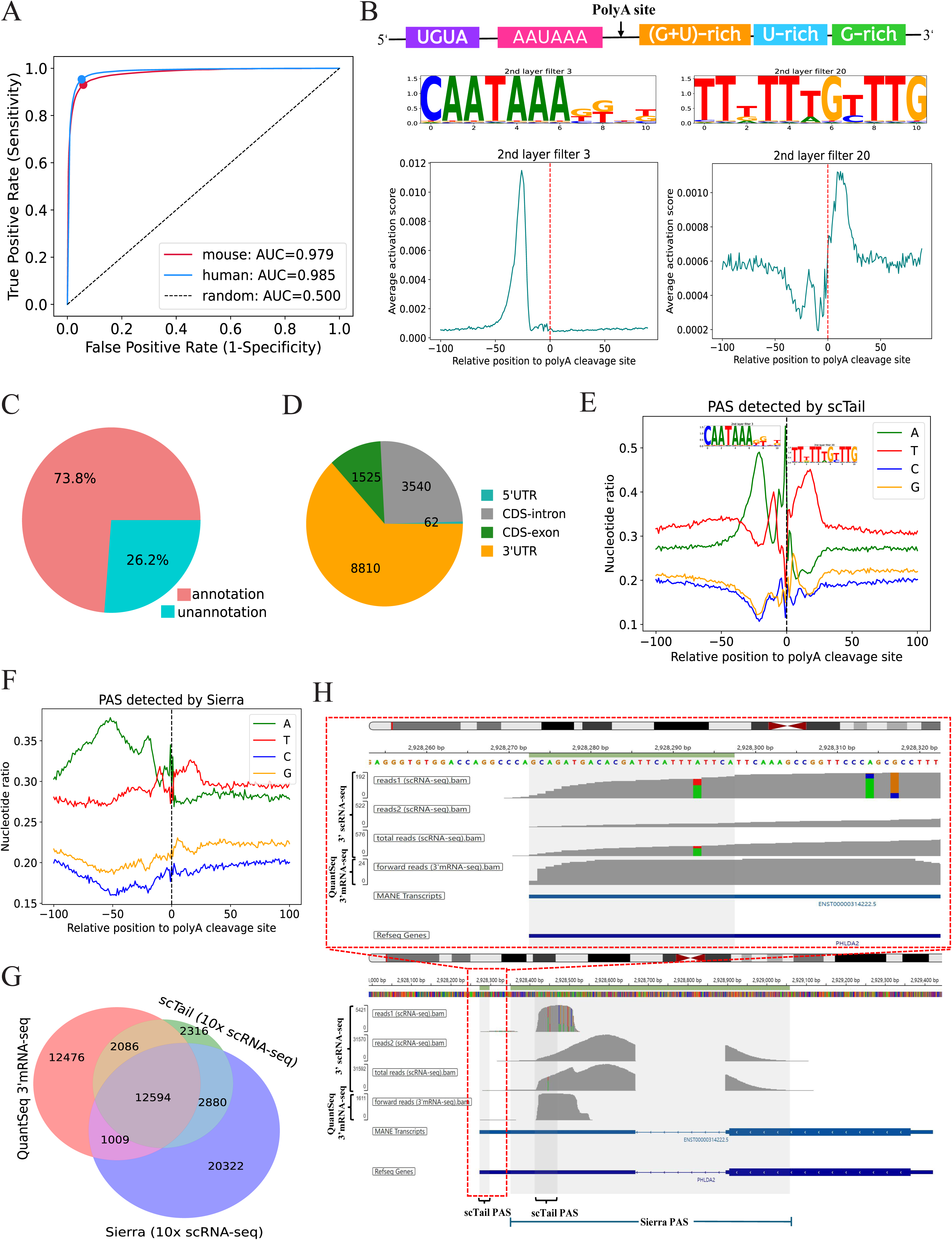
Performance of scTail detecting PAS in K562 dataset. **(A)** ROC curves for sequence model of human and mouse embedded in scTail. **(B)** Schematic showing the arrangement of motifs surrounding polyA sites (top panel). Activated motifs and their average activation score around PAs (+/- 100 bp; bottom panel). **(C)** Pie chart of the percentage of annotated and novel PASs identified by scTail in the K562 dataset. **(D)** Genomic distribution of the detected PAS. **(E)** Nucleotide distribution of sequences at PASs detected by scTail. Upstream (–) 100 bp to downstream (+) 100 bp sequences of PASs were analyzed. **(F)** Profile of nucleotide frequencies in the (+/-) 100 bp vicinity of PASs identified by Sierra. **(G)** Venn diagrams showing the overlap number of PAS from scTail, Sierra, and QuantSeq REV sequence data. **(H)** IGV track plot displaying reads coverage from reads 1, reads 2 and total reads of 10x Genomics and forward reads from QuantSeq REV for gene PHLDA2. Grey regions showing PAS detected by scTail and Sierra. The top panel is the close-up view of the red frame.

Although several methods can also detect PAS at single-cell resolution, all of them merely rely on reads2 in 10x Genomics data, hence may suffer from lack of precision and over-broad stretch. Here, we selected one widely used tool called Sierra [20] as an example to compare with scTail on the K562 dataset. Visually, the nucleotide distribution analysis supported higher accuracy of scTail than Sierra, compared to the known PAS motifs (Fig. 3F). Quantitatively, we also introduced QuantSeq REV 3’mRNA sequenceing data of K562 [31] as ground truth to evaluate scTail and Sierra. We found that scTail achieves a higher overlap proportion (73.85%) with the ground truth compared to that of Sierra (36.95%; Fig. 3G). Of the 28,165 QuantSeq REV-detected PASs (ground truth), 52.12% was also defined by scTail. In comparison, Sierra recovered fewer QuantSeq REV PASs (48.29%, Fig. 3G), despite that it returns a substantially higher number of putative PASs. One possible reason for the decreased recovery rate in the reads2-based method is the low coverage of single-cell RNA-seq, especially for the minor PAS isoform. Therefore, we performed visual inspections on example genes via the integrated genome viewer (IGV) on four tracks including reads 1, reads 2, and total reads from 10x Genomcis and also the forward reads from QuantSeq REV. As we expected, compared to the forward reads in QuantSeq REV as a technical reference, reads 1 in 10x Genomics shows a more similar distribution than its reads 2 counterparts. For PHLDA2, scTail detects two PASs, where the distal PAS is annotated and also supported by forward reads from QuantSeq REV but missed by Sierra possibly due to the proximity of the two PASs with highly imbalanced expression, suggesting an enhanced sensitivity of scTail. The proximal PAS is an unannotated PAS and identified by both scTail and Sierra, while the peak detected by Sierra is largely broader than the ones returned by scTail and forward reads from QuantSeq REV; the high consistency of the latter two also demonstrates the accuracy of scTail (Fig. 3H). We examined a considerable number of examples such as SNX11 and MAGEH1 and found that Sierra tends to return broader peaks and imprecise PAS due to the wide stretch of reads 2 and possibly nearby location of multiple PASs (Supplementary Fig. 4), which aligns with the global distribution of peak width (Supplementary Fig. 5).

### scTail facilitates cell identity and trajectory analysis

Given that scTail is able to detect PASs precisely and sensitively at a single-cell level, we wonder how much it can improve cell identity analysis and to which extent it exhibits a cell-type specificity. Here, we downloaded the transcriptomes of 14,537 epithelial cells in the human colon, ileum, and rectum from a recent study with reads1-preserved 3’ scRNA-seq data [32] to explore the variability of PAS usage across cell types and organs (Fig. 4A, B). Due to technique limitation, previous studies can only conclude that PAS is tissue-specific [33]. By utilizing scTail, we can acquire matched gene expression and PAS expression for each cell, allowing us to answer whether PAS profiles are more similar within cell types or organs. In total, 14,025 PASs were identified and quantified for all cells from 3 tissues and then used to compute a Jensen–Shannon divergence (JSD) between each cell type in each tissue (at pseudo-bulk level). When conducting hierarchical clustering on the JSD, the dendrogram reveals that samples belonging to the same cell type (across organs) exhibit a greater degree of similarity, with few exceptions (Fig. 4C). The clustering pattern characterized by the dominance of cell types implies that the majority of cell types exhibit a preserved PAS expression marker. To provide additional evidence of how the PAS profile can be used in identifying cell types, we conducted a comparison of the top 20 most prominent markers at both PAS and gene levels for each cell type. As Venn diagrams show, a partial intersection between gene- and PAS-based markers can be observed, suggesting that PAS can potentially function as supplementary features for identifying cell types (Fig. 4D). For instance, chr16 28591942 28592028*SULT1A2 (ENSG00000197165.11), chr12 27697550 27697592*REP15 (ENSG00000174236.4) and chr19 50428917 50429010*SPIB (ENSG00000269404.7) serve as a signature for enterocyte, goblet cell and paneth-like cell, respectively (Fig. 4E, Supplementary Fig. 6).

**Figure 4.**
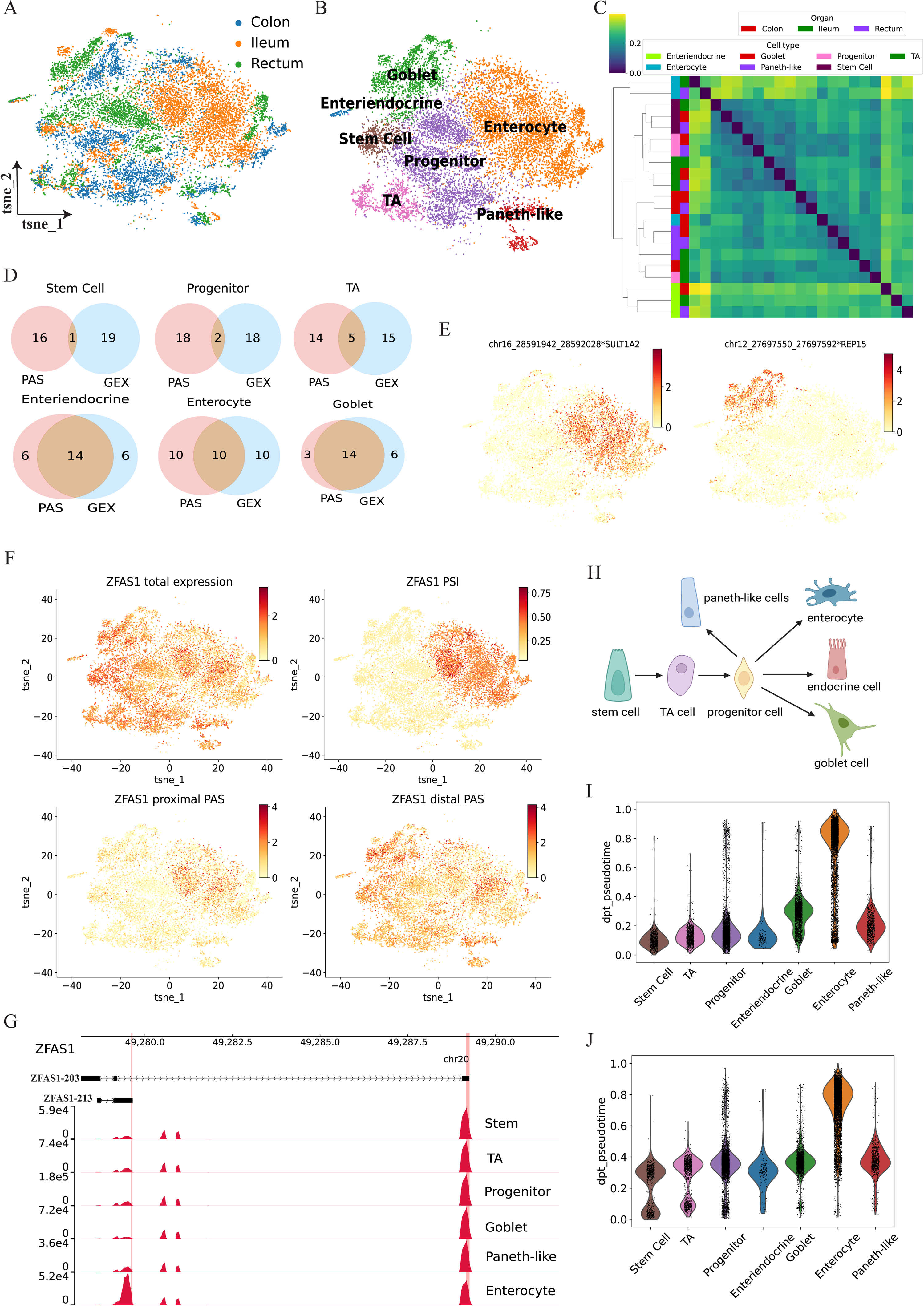
scTail assists in cell type identification and cell trajectory inference in the human intestine **(A, B)** tSNE plot of 14,537 single cells, annotated with three parts of human intestine (A) and cell type (B). **(C)** Hierarchically clustered heatmap of Jensen-Shannon divergence (JSD) of abundance of PAS among all cell types from ileum, colon, and rectum. **(D)** Venn diagrams of the top 20 significant PAS markers and gene expression markers in various cell clusters of the human intestine. **(E)** The expression of two PAS markers of enterocyte and goblet cell. **(F)** tSNE plots show total RNA expression, PSI, proximal and distal PAS expression of ZFAS1 in all human intestine cells. PSI value is estimated with cell type-specific prior. **(G)** Track plot shows read coverage of proximal and distal PAS of ZFAS1 in different cell types. **(H)** Schematic of intestine stem cell (ISC) differentiation under homeostatic conditions. **(I, J)** Violin plots of pseudotime of distinct cell types inferred by PAS expression profile (I) and gene expression profile (J).

Subsequently, our focus shifted towards genes that possess at least two PASs to analyze the alternative usage. BRIE2 was applied to search for genes with differential alternative PAS usage for each cell type compared with the rest of all other cell types. Note, for genes with more than two PASs, only proximal PAS (the closest PAS to the transcription start site) and distal PAS (the furthest PAS to the transcription start site) were utilized. In total, 2,189 genes with significant differential alternative PAS usage (FDR < 0.05) were found in all cell types (Supplementary Fig. 7A-G, Supplementary Table. 1). The volcano plot reveals a pronounced asymmetrical trend of significant genes, which hints that several cell type prefers to apply proximal PAS, such as TA and enterocyte whereas stem cell, progenitor, goblet cell and paneth-like cell would like to make use of distal PAS (Supplementary Fig. 7H). Among those genes with significant differential PAS usage in a certain cell type, we discovered several genes with significant shifts in PAS abundances in one cell type while the gene-level expression remains consistent across all cell types. Fig. 4F shows an example of such a gene ZFAS1, which is known as a protein regulator involved in colorectal cancer tumorigenesis and development [34]. Although there is no alteration in the gene expression of ZFAS1 across these cell types, we observed that the proximal PAS of the gene is more highly expressed in enterocytes compared to other cell types. The ZFAS1 gene undergoes an isoform shift in the enterocyte, where the proportion expression of the proximal PAS (e.g. PSI) is significantly higher, indicating distinct cellular PAS localization (Fig. 4F), with high concordance in the track plots (Fig. 4G). Another example gene is NEDD4L, which mediates Wnt3 ubiquitination and modulates gut microbiota to control colorectal cancer progression [35]. The distal PAS of NEDD4L marks the goblet cell despite constant gene abundance (Supplementary Fig. 8). Taken together, these findings demonstrate that PAS-level abundance can assist in refining cell types beyond what is possible blind at gene-level expression.

During the interstinal differentiation, the stem cells sequentially differentiate into transit-amplifying (TA) cells and progenitor cells. Then, the secretory progenitor cells migrate upwards to become goblet cells, enteroendocrine cells, and paneth cells, while the absorptive progenitor cells divide to enterocyte [36] (Fig. 4H). To determine the role of PAS in deciding cell fate, the PAS-level and gene-level expression profiles (for genes with significant alternative PAS usage) were exploited to infer the pseudotime of cells through geodesic distance along the graph (Supplementary Fig. 9). As shown in Fig. 4I and 4J, the pseudotime inferred from both PAS and gene profiles gradually increases from stem cell to TA and to progenitor. Also, the pseudotime inferred from the PAS expression profile of goblet, enterocyte, and paneth-like cells is higher than progenitor cell, which is consistent with the process of differentiation in the human intestine (Fig. 4I). However, the pseudotime inferred from gene-level expression is almost the same for progenitor, goblet and the paneth-like cells (Fig. 4J). We suspect that the PAS expression profile provides extra information than the gene expression profile possibly due to alternative PAS usage. This finding is consistent with a previous report that instead of alterations in gene expression, pre-mRNA processing including splicing and polyadenylation is responsible for reshaping the transcriptome and proteome during the loss of stemness and lineage commitment [37].

### Altered PAS usages in esophageal squamous cell carcinoma micro-environment

Although comprehensive analyses of alternative polyadenylation (APA) usage have been conducted in various cancer types at bulk level [6, 10], the study exploring APA usage at single-cell resolution was limited to reads 2, lacking accuracy of detecting PAS [38]. Here, scTail was utilized to an esophageal squamous cell carcinoma (ESCC) dataset (3’ scRNA-seq, 10x Genomics) of both adjacent nonmalignant and matched tumor samples from 11 ESCC patients, covering 89,313 cells in total [39] (Fig. 5A, B, Supplementary Fig. 10) to delve into what is the profile of APA usage across various cell types in ESCC and regulatory mechanism behind it. With scTail, we identified 9,888 PASs, where 1,831 genes contained at least two PASs (Supplementary Fig. 11). By utilizing BRIE2 [40] on these multi-PAS genes, a grand total of 1,005 genes exhibited significant divergence in the usage of alternative PAS between normal and tumor in any cell type (FDR < 0.01; Supplementary table 2). It should be noted that we only used the proximal and distal PASs if there are multiple (definition mentioned above) as input for BRIE2, and as we expected, where distal PASs have a higher proportion overlapped with 3’ UTR compared with that of proximal PAS (Supplementary Fig. 12). Take epithelial cell as an example, among genes exhibiting alternative PAS activation in tumor, several are widely recognized as cancer-related biomarkers such as S100A14 serving as modulator of terminal differentiation in esophageal cancer [41], PITX1 relating with RAS Activity and Tumorigenicity [42] (Fig. 5C). In addition, notably differential genes with APA usage between normal and tumor conditions can be identified by BRIE2 for each cell type (Supplementary Fig. 13). To further assess the usability of these genes, the proportion of proximal PASs (i.e., PSI) of genes, the gene expression and combination of PSI and gene expression were feed to logistic regression to predict binary disease status with tenfold cross-validation separately. All three feature groups achieved good predictions on cell types that are closely relevant to cancer, e.g., epithelial cells (Fig. 5D; Supplementary Fig. 14; only tested on cell types with >500 cells). Impressively, combining the extra PSI matrix to the gene-level expression consistently improves the prediction performance in terms of AUROC, not only for those moderately predicted cell types, e.g., B cells but also those more accurately cell types, e.g., fibroblast cells (Fig. 5D). In general, among these significant differential genes, the number of genes preferring to utilize proximal genes is obviously more than the number of genes favoring expressing the distal PAS in tumor samples (Fig. 5E), indicating global shorting in tumor condition and aligning with the previous reports [6, 10].

**Figure 5.**
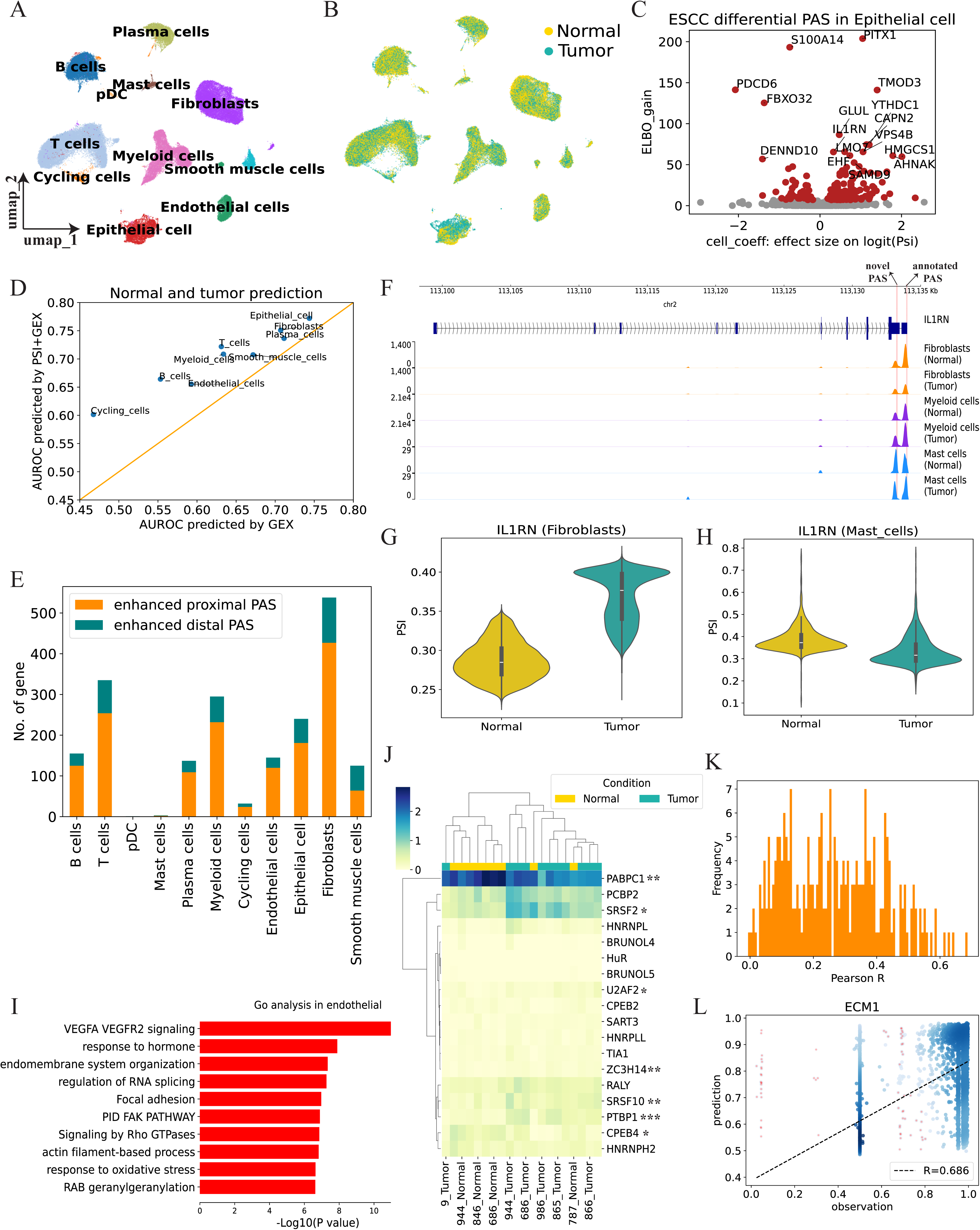
scTail detected polyadenylation site switch from esophageal squamous cell carcinoma. **(A, B)** UMAP visualization of all cells (n=89,313) in esophageal squamous cell carcinoma. Each dot represents an individual cell, where colors indicate cell type (A) and disease status (B). **(C)** Volcano plot illustrating the correlation between ELBO_gain and effect size on logit(PSI) for detecting differential PAS between normal and tumor cells. The effect size on logit(PSI) is represented by cell_coeff. A positive value means higher PSI in normal cells. ELBO_gain reflects the evidence lower bound difference for the two hypotheses (Methods). **(D)** Scatter plot showing AUROC predicted by expression only and expression plus PSI of significant genes with APA usage by exploiting logistic regression model. Tenfold cross-validation was utilized to evaluate the model. **(E)** Bar plot exhibits number of genes preferring to use proximal or distal PAS in tumor condition for distinct cell types. **(F)** Genome track plot of IL1RN in fibroblast, myeloid cell and mast cell of normal and tumor. A single horizontal genome track denotes the coverage across all cells within a specific cell type. The red line indicates the location of PAS detected by scTail. **(G, H)** Violin plot on example gene IL1RN for fibroblast (G; n = 9071 cells for normal; n = 8450 cells for tumor) and mast cell (H; n = 757 cells for normal; n = 603 cells for tumor). The y-axis PSI represents the proportion of proximal PAS among the abundance of total PAS in each cell type. **(I)** Bar plot displaying the enriched GO terms of genes preferring to use distal PAS in tumor condition in endothelial cells. **(J)** Heatmap shows the hierarchical clustering of patients by the abundance of RBPS that binds with gap sequence of significant differential genes with APA between normal and tumor. ***: FDR < 0.0001, **: FDR < 0.01, *: FDR < 0.05. **(K)** Histogram of Pearson’s correlation between predicted and measured expression of significant differential genes with APA in epithelial cells. Prediction was conducted by random forest. Tenfold cross-validation evaluates the performance of the model. **(L)** A scatter plot between the predicted with all RBP abundance and measured expression of ECM1 in epithelial.

Additionally, we also observed cell-type specific PAS usage in the ESCC dataset, such as IL1RN, which is known to be downregulated in ESCC patients [43]. As shown in Fig. 5F, scTail identified two cell-type specific PASs, including proximal PAS (novel PAS) and distal PAS (annotated PAS). Interestingly, if we focused on the proportion of two detected PASs in fibroblast, we found that the proportion of the proximal PAS shows a significant up-regulation in tumor condition compared to the normal condition due to the decreased abundance of distal PAS in tumor (Fig. 5G, Fig. 5F). However, mast cell exhibits a reverse trend, where the proportion of proximal PAS decreases in tumors because of decrease expression of proximal PAS in tumor (Fig. 5H, Fig. 5F). Surprisingly, the proportion of proximal PAS did not change a lot between normal and tumor in myeloid cells (Fig. 5F, Supplementary Fig. 15), highlighting the intricate nature of PAS regulation and its concurrent influence on cellular phenotypes and pathological states. To further characterize the functions associated with APA in ESCC, we conducted a pathway enrichment analysis using genes with normal- or tumor-specific PASs at distinct cell type levels. Specifically, in endothelial, those mRNA transcripts applying proximal PAS exhibit enrichment in GO terms relevant to VEGFA VEGFR2 signaling which is known to be correlated with the prognosis of patients with ESCC [44], response to hormone and endomembrane system organization, collectively indicating that APA plays an essential role in the molecular features of ESCC (Fig. 5I). Besides endothelial cells, transcripts preferring to proximal or distal PAS in other cell types also contribute to the progress of ESCC (Supplementary Fig. 16).

To elucidate the potential regulatory mechanism underlying the global preference of proximal PAS, we searched for motifs appearing in the gap sequence (i.e. sequence from proximal PAS to distal PAS) of significant differential genes with APA usage by using FIMO (v4.11.2) and further detected the RBPs binding with these motifs (Supplementary Fig. 17). Here, we took Polyadenylate-binding protein cytoplasmic 1 (PABPC1) as an example and surveyed the CISBP-RNA database [45], and found three binding motifs: M146 0.6, M275 0.6 and M349 0.6. M146 0.6 closely resembles the sequence logo detected by FIMO in the gap sequence, which implies potential regulatory elements of these shortened sequences in the ESCC tumor (Supplementary Fig. 18). Unbiased hierarchical clustering analysis of these RBPs expression effectively segregated tumor and normal samples, indicating that they may play a critical role in the regulation of alternative polyadenylation usage in ESCC (Fig. 5J). For example, PABPC1 was downregulated in the tumor (FDR=0.002; Supplementary Fig. 19), which is consistent with previous studies [46, 47]. Due to the characteristics of PABPC1 (i.e., polyadenylate-binding), the downregulation of its expression and the shorter 3’UTR in the tumor are harmonious.

We also tried to discover essential RBPs from other angles. Expression profiles of 2,960 annotated human RBP were provided to a random forest regressor to predict PSI of 362 genes that were detected with alternative polyadenylation usage in epithelial cells. As Fig. 5K shows, the PSI values of several genes can be predicted well from RBP expression, such as ECM1 (Pearson’s R = 0.686), RAN (Pearson’s R = 0.639), S100A2 (Pearson’s R = 0.630) and TIMM13 (Pearson’s R = 0.620) (Fig. 5L, Supplementary Fig. 20). We further studied how expression of each RBP contributes to the prediction of these four genes. Interestingly, most of the expressions of RBPs have a positive correlation with the PSI of ECM1, RAN, and S100A2, and a negative correlation with the PSI of TIMM13 (Supplementary Fig. 21). We suspect that the main contributing factor is the preference for TIMM13 to utilize distal PAS (average PSI = 0.32), while the other three genes tend to favor the usage of proximal PAS (RAN: average PSI = 0.74; S100A2: average PSI = 0.76; ECM1: average PSI = 0.70). To examine the function of these top features, Gene ontology (GO) analysis using 33 RBPs revealed significant enrichment under several major GO categories, including ribosome, cytoplasmic, stomach neoplasms, and neutrophil degranulation (Supplementary Fig. 22), relevant with alternative polyadenylation usage and characteristic of the dataset. The mRNA 3’-end formation is a co-transcriptional process that is mediated by the cleavage and polyadenylation (CPA) machinery, which consists of a multiprotein core complex and associated factors (i.e. CPA specificity factor CPSF) [1]. Based on this, we asked which CPSF contributes to the PAS shift of ESCC. Expressions of CPSF between normal and tumor were compared and we found that CPSF3 and SCAF8 exhibit significant differences (Supplementary Fig. 23). SCAF8 has been reported as an anti-terminator protein, which is consistent with its lower expression and global proximal PAS usage at tumor condition [48].

### Polyadenylation site shifting during mouse forelimb histogenesis

Lastly, we aimed to examine the contributions of alternative PAS usage in mouse development. Here, as a showcase, we focused on forelimb development during the embryonic stages by analyzing the data generated from the ENCODE Consortium mouse embryo project [49]. In this dataset, the forelimb includes 70,376 cells annotated into 25 cell types involved in 7 samples from embryonic day 10.5 (E10.5) to E15.0 (Fig. 6A, B). In total, 24,109 PASs in 12,974 genes were detected by scTail (Supplementary Fig. 24). To analyze the PAS switch, we focused on genes (n = 5,849) with multiple PASs and utilized BRIE2 to inspect their PAS usage between each time point and the rest other times at cell type level. We identified 2,804 and 10,619 significant differential genes with PAS shift in muscle and mesenchymal for all time points (FDR < 0.01; Supplementary Table 3). As shown in Fig. 6C and Supplementary Fig. 25, several genes with notable alternative polyadenylation site usage contribute greatly to organogenesis in muscle or mesenchymal, such as Tpm2 associated with embryonic skeletal muscles [50], Ddx18 associated with mESCs differentiation [51], Hes1 contributing to differentiation responses of embryonic stem cells in mouse [52]. As an example, muscle cells dominantly expressed the distal PAS of Tpm2 at E10.5 and E11.0 but switched to its proximal PAS from E12.0. Also, mesenchymal cells preferred to express proximal PAS of Pim1 at E10.5, while shifting to distal PAS from E11.0 (Fig. 6D). We observed that the change of PSI in individual cells is consistent with time-dependent APA shift of Tpm2 and Pim1 (Fig. 6E). However, the gene-level expression of these two genes is broadly consistent without any significant difference along the time axis (Fig. 6F). This further highlights the complementary function of APA to bolster the ability to distinguish cell states in scRNA-seq data. Furthermore, with functional enrichment analysis, we found the genes with significant PAS shift between E10.5 and rest time points in muscle cells mainly affected the cell division and molecular catabolic process including the metabolism of RNA, protein catabolic, and protein-DNA complex organization (Fig. 6G). In addition, we also compared the expression of CPSF among different embryo stages and discovered that only Cpsf7 gradually increased over time and significantly escalated from E12.0 (Supplementary Fig. 26; Supplementary Fig. 27). This is consistent with the finding that changed abundance of CPSF7 (one component of CFIm) is responsible for the switching of UTR lengths in the development of mammalian early embryos [53].

**Figure 6.**
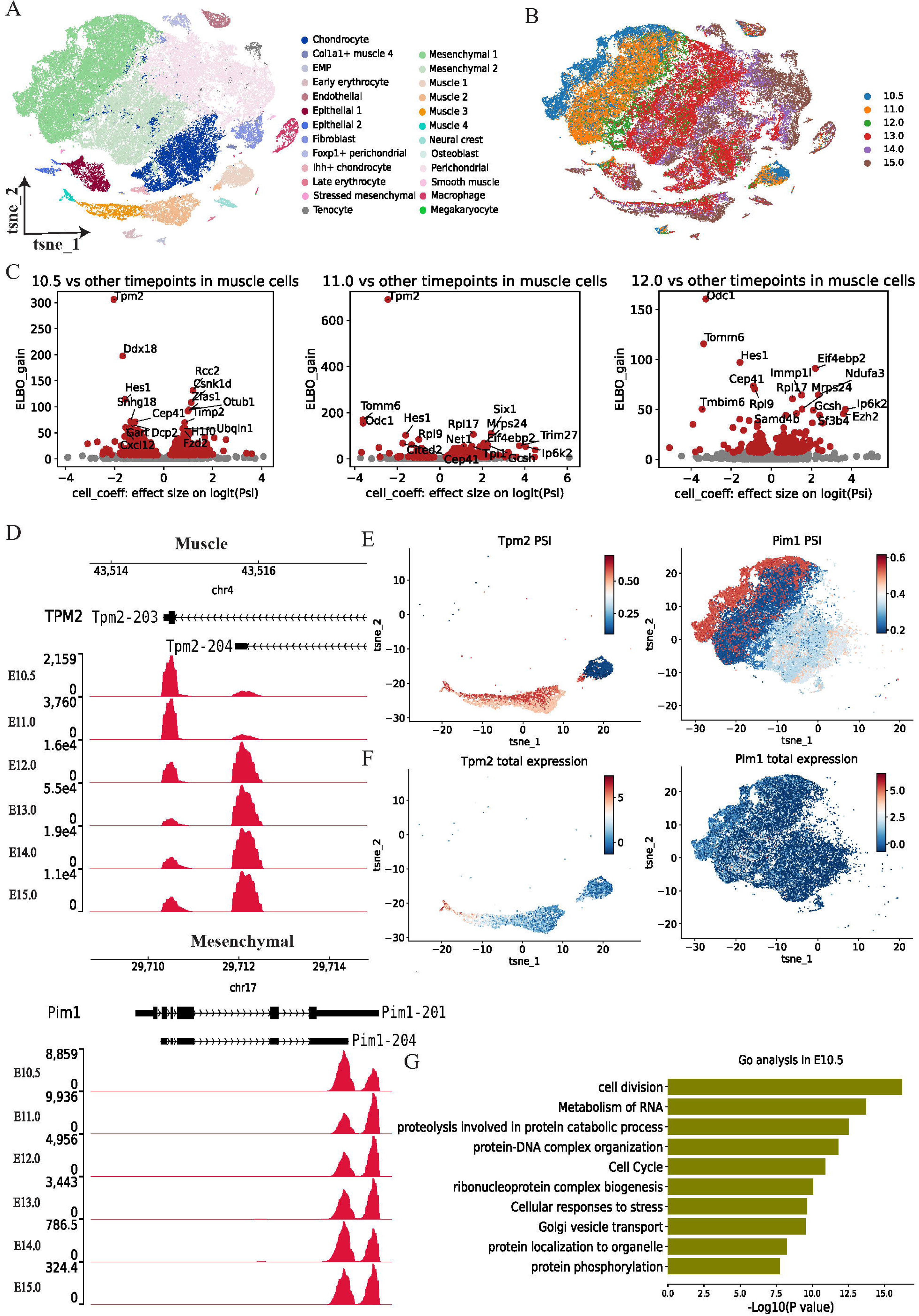
scTail identifies differential alternative PAS usage from mouse forelimb development. **(A, B)** tSNE visualization of all cells (70,376) in forelimb histogenesis. Each dot is the individual cell, with colors coded according to the cell types (A) and development stages (B). **(C)** Volcano plot showing the correlation between ELBO_gain and effect size on logit(PSI) for identifying differential PAS between E10.5 and other time points (left), E11.0 and other time points (middle), E12.0 and other time points (Bottom). Cell_coeff represents the effect size on logit(PSI). Positive value indicates higher PSI in E10.5 (left), E11.0 (middel) and E12.0 (right), respectively. ELBO_gain marks the evidence lower bound difference for the two hypotheses (Methods). **(D)** Genome track plots display read coverage of Tpm2 and Pim1 in various development stages of muscle cell (top) and mesenchymal cell (bottoem). **(E)** tsne plots visualize the mapping of PSI value of Tpm2 (left) and Pim1 (right) on individual cells of muscle and mesenchymal. PSI value is estimated with stage-specific prior. **(F)** tsne plots display the mapping of the abundance of Tpm2 (left) and Pim1 (right) on individual cells of muscle and mesenchymal. **(G)** Bar plot showing the enriched terms based on genes with significant differential PAS usage between E10.5 and other stages in muscle cell.

## Discussion

In this study, we present a computational method called scTail that can identify PASs by exploiting paired reads 1 (length > 100bp) and quantify PAS by using total reads 2 from 3’ scRNA-seq data (10X Genomics). Our method, scTail, is designed in a data-driven manner and incorporates a sequence model that identifies PASs accurately. This enables us to efficiently detect alternative TSS usage between different single-cell populations by utilizing the BRIE2 model seamlessly. Following that, we focused on the analysis of alternative PAS usage in various biology scenarios including human intestinal differentiation, esophageal squamous cell carcinoma conditions, and mouse forelimb histogenesis, where using the right PAS is an important element of the regulation, presumably relating to RBP and CPSF. Therefore, PAS expression usually offers supplementary information to the gene-level expression. Overall, the easy-to-access PAS analyses bring new opportunities to reveal biological insights while along with multiple challenges to be further addressed.

From the technology perspective, multiple techniques with bulk sequencing have been introduced for a while, such as PolyA-seq [12] and QuantSeq REV [15], allowing us to annotate the PAS in common conditions. These technologies utilized novel protocols to directly target the 3’ end of polyadenylated transcripts, where the start nucleotide of the read-out corresponds to the authentic transcription end site. However, similar technologies at single-cell resolution are relatively scarce, and the single-cell polyadenylation sequencing (scPolyA-seq) [54] is the only example highlighting the capability in analyzing PAS usage. On the other hand, multiple common scRNA-seq protocols are 3’ biased and poly-A selected, therefore in principle can capture the PAS. Recently, Fansler and colleagues showed that the Microwell-seq platform has a relatively precise capture of the PAS, hence can measure the 3’ UTR length in single cells [55]. Nevertheless, we are still far from making PAS analysis a routine with easy access. With our scTail, we demonstrated that the mainstream platform 10x Genomics is nearly perfect for PAS analysis, as long as the cDNAs in reads 1 are sequenced and preserved. Still, more efforts are needed to emphasize this message, ideally having this functionality included in the default processing pipeline.

In terms of biology, the 3’ UTR has profound impacts on a wide range of biological functions, and the three case studies we presented are only the tip of the iceberg. As we evidenced, there are prevalent associations between alternative PAS usage and development and diseases. Still, the regulatory mechanism remains largely unknown, for example, how multiple RNA binding proteins work collectively to modulate the mRNA functions. This closely links to the community’s attention to the genetic regulation of the switch of 3’ UTR [56], some of which show strong colocalization with disease variants, e.g., ZC3HAV1 in multiple sclerosis [57].

Regarding therapeutics, mRNA is increasingly popular as a vaccine and drug, while how to design and optimize 3’ UTR sequences is only an emerging challenge and opportunity [58]. Broadly speaking, precise detection of PAS in a diverse context may allow us to compile a large amount of training set for learning the broad language of the 3’ UTR sequence, which can facilitate resolving specific tasks in 3’ UTR designs for different properties.

## Methods

### scRNA-seq initial data analysis

Raw fastq files contains reads1 (> 100 bp) and reads2 were downloaded, and then reads were aligned to the *homo sapiens* reference genome (hg38) or *Mus musculus* reference genome (GRCm39) to produce pair-end alignment bam file by utilizing STAR [59] (v2.7.11) (parameter: `--soloType CB UMI Simple --soloStrand Reverse --clip5pNbases 60 0 --outFilterScoreMinOverLread 0.1 --outFilterMatchNminOverLread 0.1`). Of note, for `--readFilesIn`, we set the order of input reads as reads1 (with cell barcode and UMI) and reads2 and this order corresponds with `--soloStrand Reverse`. Then we filtered reads whose ‘GX’, ‘CB’, and ‘UB’ tags are ‘-’ in the aligned bam file. Sometimes, the bam file of each sample is too large to run scTail fast, and Sinto (v 0.10.0) was utilized to split one bam file into two or more smaller bam files randomly.

### Implementation of scTail

scTail is a computational method to detect PAS and it consists of two major stages: identify PAS by using long reads 1, optionally merge PASs from individual samples, and assign reads 2 to the region of PAS.

### Step 1: identification of PAS

Pysam [60] was used to parse the preprocessed bam file to obtain UMI counts of each reads end (Fig. 2A). Next we exploited paraclu [28] to perform clustering and filter clusters according to the thresholds set by users. The thresholds required to set include maximum cluster length, minimum density increase, and minimum cluster count. Except for these filter parameters, an embedded convolutional neural network also helps with filtering.

The deep neural network only supports two species: human and mouse. Positive samples of human PAS (n = 251,072) came from stringent PASs dataset which includes PolyA DB3, PolyA-seq, and PolyASite or GENCODE annotation in previous study [24]. Positive samples of mouse PAS (n = 106,894) are the sum of GENCODE annotation of the protein-coding gene (n = 21,786) and PAS dataset (n = 85,108) with high confident PAS from previous study [24]. Negative samples of human (n = 13,809) and mouse (n=92,164) were randomly selected from intergenic regions without overlapping with any positive samples. To keep positive and negative samples balanced, some transcription start sites were randomly selected as additional negative samples. Finally, 251,072 human negative samples and 106,894 mouse negative samples were obtained.

Sequences of upstream 100 bp and downstream 100 bp of positive or negative PAS were extracted by the python package pyfaidx [61] from the fasta file. Then we utilized kipoiseq [62] package to convert DNA sequence to one-hot encoding (A = [1,0,0,0], C = [0,1,0,0], T = [0,0,1,0], G = [0,0,0,1], N = [0,0,0,0]). PyTorch (v2.1.0) was used to construct the convolutional neural network, which is composed of three convolution layers followed by a Rectified Linear Unit (ReLU) for activation, connected by batch normalization, max pooling, and dropout layer with dropout rate 0.4. The flattened output from the dropout layer was passed through fully connected layers (32 neurons) with a ReLU activation function. Next, the flattened output was connected to a second fully connected layer consisting of 2 neurons that utilized the sigmoid function to determine the classification probability. The initial convolutional layer consists of 128 filters, each with a width of 8-mer and 4 channels. The subsequent convolutional layer consists of 32 filters, with each filter having a size of 4x4. The third convolutional layer has 64 filters, where the filter size is 2x2. This model can be represented in the following formula:

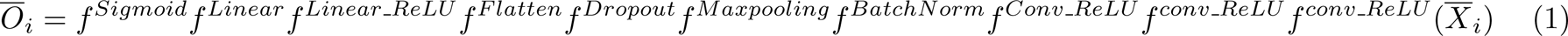

where 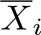 represents the one-hot encoded matrix of size 4 by 200. The total samples were split into three sets: train, test, and validation using a ratio of 6:2:2. The model was trained using a batch size of 128 and 100 epochs, employing the Stochastic Gradient Descent (SGD) optimizer with a learning rate of 0.003 and momentum of 0.8. The model embedded in scTail is the one with the lowest loss evaluated by the validation set.

When there are multiple samples, scTail aims to compile a union set of PAS across samples. Specifically, in the last step above, we can detect PAS for individual samples. Then, the bedtools was applied to merge PAS from all samples (parameter: `-d 40 -c 4 -o collapse`).

### Step 2: quantification of PAS

Here, we introduced a quantification model to assign reads 2 to the merged PAS. The distance between the termination of reads 1 and reads 2 was defined as fragment size. In this model, it is assumed that fragment size follows a univariate log-normal distribution:

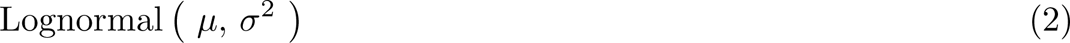

Assuming *x_i_*is the fragment size for assigning the *i*th read 2 to a certain PAS (namely *x_i_* denotes the distance between the read2 end and the PAS), the likelihood can be estimated by:

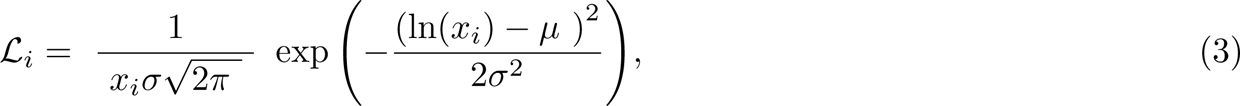

where *µ* and *σ* are the parameters of the lognormal distribution and the likelihood was calculated by using the scipy python package (v1. 12.0; log-normal distribution). Here, we estimate these two parameters by using the empirical distribution of fragment size of the paired reads 1 and 2 only from genes with a single PAS. Specifically, the fragment of each read pair is calculated by the distance between the end positions of read 2 were obtained by parsing the aligned bam file via pysam and that of its paired read 1. Then, we used a heuristic way to estimate the parameters by transforming the mean *µ_X_* and the standard deviation *σ_X_*of the fragment sizes, as follows,

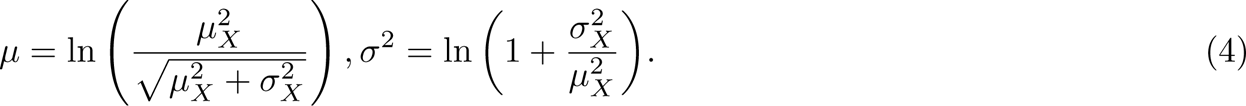

Next, we decided to keep or filter this query read by comparing this likelihood in Eq. 3 and the threshold defined by the half of likelihood of emperical distribution of fragment size. If there are multiple PASs that exist in a gene, we assign the read to the PAS with the highest likelihood that passes the threshold. Finally, we summed the total reads of each PAS.

Of note, in both PAS identification and quantification, we all first eliminate PCR duplicates and save only one read per UMI in the same cell. The reads with the furthest distance to the transcription start site were selected when reads from the same UMI and the same cell aligned to different positions.

### Analysis of genomic feature of PAS

The genomic interval of 5’ UTR, 3’ UTR, intron, and exon were obtained by inputting the hg38 GTF file to *gencode regions* package (https://github.com/saketkc/gencode regions). Then PAS detected by scTail in K562 were assigned to genomic features such as 5’ UTR, 3’ UTR, and so on.

### Explanation of convolutional neural network

The hidden features within the convolution neuron encode local sequence combinations that can be leveraged to accurately predict the PAS. The method called Maximum Activation Seqlet [30] was utilized in this study. By utilizing this method, it becomes possible to identify the brief sequence segments that the convolutional filters are detecting. These segments can then be employed to create Position Weight Matrices (PWMs). In this approach, we initially divided the input sequence X into multiple subsequences (Seqlet) 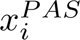 that have the same length as the receptive field of the target neuron *c*. We searched for Seqlet from X that have the highest activation values for convolutional filter *c* and bagged them into *P_c_*.

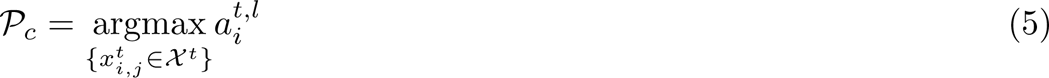

By stacking the short sequences in *P_c_* together, we can determine the frequency of each nucleotide at every position, resulting in the creation of Position Weight Matrices (PWMs). Afterward, the convolutional filters were transformed into PWMs and were then visualized as sequence logos using a Python package logomaker (v 0.8).

### QuantSeq 3’mRNA-seq preprocessing

Raw fastq file of the K562 dataset (QuantSeq 3’mRNA-seq) was downloaded from the NCBI SRA database. Next, we applied cutadapt (v4.6) to remove 19 bases (i.e. PolydT) from reads 1 (parameter: `-u 19`). Then STAR was employed to align the clean forward reads to the human reference genome (hg38) with the default parameter.

### Preprocessing of all dataset

For one dataset, we can get two cell-by-feature UMI matrices, both in h5ad format (i.e. PAS expression h5ad file and gene expression h5ad file). PAS and gene expression matrices were obtained respectively from the output of scTail and the combination of the output of STAR aligner. Cell type annotation and the 2D coordinates of t-SNE or UMAP were obtained from the dataset source and mapped to these two matrices. Scanpy (v.1.9.5) [63] was utilized to perform normalization by total counts (target sum=1e4) and extracted highly variable genes.

### Identification of alternative polyadenylation site usage on cell type, disease and development stage

BRIE2 (v 2.2.2) [40] was employed to detect polyadenylation site shift across different cell types, disease status, and development stages. In the case of ESCC and mouse forelimb, we constructed an h5ad file with two layers containing the abundance of two PAS (raw UMI count) for each cell type. Of note, for genes with multiple PASs, the closest and farthest PAS to the transcription start site were selected, as proximal and distal PASs. The design matrix comprises cell detection rate and cell state information (for example: tumor: 1 and normal: 0; E10.5: 1 and other development stages: 0) for each cell type. In the dataset of human intestinal, the h5ad file includes all cell types and the design matrix consists of cell detection rate and cell type information. The brie-quant module was utilized to identify APA for all pairwise comparisons (parameter: `--batchSize 1000000 --minCell 10 --interceptMode gene --testBase full --LRTindex 0` and FDR *<* 0.01 was defined to select genes with significant differential PAS usage.

In addition, when we show the PSI value of individual cells, the prior was changed. In the human intestinal dataset, brie-quant was run using all cell types as prior; namely the model will learn a separate prior for each cell type (Fig. 4F). In the case of mouse forelimb, all stages were utilized as prior; namely the model will learn a separate prior for each stage (Fig. 6E).

### Hierarchical clustering analysis

To investigate the similarity of PAS profiles across three tissues (i.e. colon, ileum, and rectum) and cell types, we exploited normalized PAS to get expression of each cell type of each tissue by using the pseudobulk method. Next, `scipy.spatial.distance.jensenshannon` was utilized to compute the Jensen-Shannon distance (JSD) between two cell types. Finally, the distance matrix was used to conduct a hierarchical clustering analysis.

### Diffusion pseudotime analysis

We first chose genes with significant differential alternative polyadenylation sites in all cell types in the human intestinal dataset and then did preprocessing in these PAS and gene expression matrices. One of the stem cells was defined as root cell and then scanpy was applied to calculate diffusion pseudotime with the function `scanpy.tl.diffmap`. Next, we used scatter and violin plots to visual the estimated pseudotime.

### Disease states prediction

To perform the disease prediction in the ESCC data, we focused on cell types that have more than 500 cells. Within each of these cell types, genes with significant differential APA between normal and tumor (FDR *<* 0.01) were selected for the analyses. PSI values of these genes were fed to logistic regression to predict disease conditions. Tenfold cross-validation was utilized to evaluate the performance of the prediction.

### Track plot visualization

IGV-Web was applied to plot reads 1, reads 2, total reads, and the forward reads from QuantSeq REV in the K562 dataset. For track plots in human intestinal, ESCC and mouse forelimb datasets, sinto was utilized to subset the bam file for each track according to their cell barcode list, such as for a certain cell type. Next, these bam files were input to pyGenomeTracks (v.3.8) [64] to visualize.

### Motif binding frequency analysis

In the ESCC dataset, we focused on genes with significant differential alternative PAS between normal and tumor conditions. For genes with multiple PASs, only the closest and furthest PASs were selected for analysis, denoted as proximal and distal PASs, respectively. Gap sequences between them were extracted by seqtk (https://github.com/lh3/seqtk). Next, FIMO (v 4.11.2 MEME-suite) [65] was used to search RBP motif occurrences within the gap sequences according to meme file of homo sapiens RNA-binding motifs PWM [45].

### proportion of proximal PAS prediction in ESCC dataset

A total of 2,961 RNA-binding proteins (RBPs) from homo sapiens were obtained from the EuRBPDB database [66]. Normalized abundance of these RBPs was fed to a random forest regressor (implemented with scikit-learn v1.3.2) to predict the proportion of proximal PAS (i.e. PSI) of genes with significant differential PAS usage in the disease condition. Pearson’s correlation was calculated between measured PSI and predicted PSI. The random forest’s function `feature importances` was utilized to calculate feature importance.

### Functional enrichment

The Metascape online web server (v3.5.20230101) [67] [options: Expression Analysis] was exploited to perform GO enrichment analysis and the top 20 enriched terms were selected to do enrichment network visualization.

## Supporting information

This file includes all supplementary figures.

## Data availability

For K562 data, its 10x Genomics 3’ scRNA-seq [68] and QuantSeq 3’ mRNA-seq [31] were downloaded respectively from GEO under the accessions “GSE231382” and National Center for Biotechnology Information (NCBI) BioProject database under the accessions “PRJNA794041”. The previously published 3’ scRNA-seq data that were reanalyzed in this study are available in the NCBI BioProject database under accession number “PRJNA777911” (ESCC dataset [39]), GEO under accession number “GSE125970” (human intestinal dataset [32]) and ENCODE under accession number “ENCSR713GIS” (mouse forelimb dataset [49]).

## Codes availability

scTail is an open-source Python package publicly available at https://github.com/StatBiomed/scTail. Detailed documentation and analysis procedures to reproduce results in this paper are also uploaded to this repository.

## Author contributions

Y.H. conceived and supervised this study. R.H. implemented the scTail and performed all data analysis. R.H. and Y.H. wrote the manuscript.

## Competing interest statement

The authors declare no competing interests.

## Acknowledgments

We thank Weizhong Zheng for technical help on the CNN model, Austin E. Gillen, Rui Fu and Xianjie Huang for reads 1 exploration, Lingyu Li and Chen Qiao for troubleshooting, and Peter Chan for server support. We also thank researchers for sharing their processed data and annotations: Li-Yan Xu and Feng Pan from Shantou University (ESCC dataset), Ye-Guang Chen and Wanlu Song from Tsinghua University (human intestinal dataset & discussing our results), Hashem Koohy and Agne Antanaviciute from the University of Oxford. This project is supported by the National Natural Science Foundation of China (No. 62222217), Innovation Technology Commission Funding (Health@InnoHK), and the University of Hong Kong through a startup fund and a seed fund (Y.H.). R.H. is supported by the Postgraduate Scholarship of the University of Hong Kong.

## Notes

### Competing Interest Statement

The authors have declared no competing interest.

## References

1. Mitschka, S. & Mayr, C. Context-specific regulation and function of mRNA alternative polyadenylation. Nature Reviews Molecular Cell Biology 23, 779–796 (2022).

2. Stroup, E. K. & Ji, Z. Deep learning of human polyadenylation sites at nucleotide resolution reveals molecular determinants of site usage and relevance in disease. Nature communications 14, 7378 (2023).

3. Lima, S. A. et al. Short poly (A) tails are a conserved feature of highly expressed genes. Nature structural & molecular biology 24, 1057–1063 (2017).

4. Agarwal, V., Lopez-Darwin, S., Kelley, D. R. & Shendure, J. The landscape of alternative polyadenylation in single cells of the developing mouse embryo. Nature communications 12, 5101 (2021).

5. Sommerkamp, P. et al. Differential alternative polyadenylation landscapes mediate hematopoietic stem cell activation and regulate glutamine metabolism. Cell stem cell 26, 722–738 (2020).

6. Zingone, A. et al. A comprehensive map of alternative polyadenylation in African American and European American lung cancer patients. Nature communications 12, 5605 (2021).

7. Mayr, C. & Bartel, D. P. Widespread shortening of 3’ UTRs by alternative cleavage and polyadenylation activates oncogenes in cancer cells. Cell 138, 673–684 (2009).

8. Dorrity, T. J. et al. Long 3’ UTRs predispose neurons to inflammation by promoting immunostimulatory double-stranded RNA formation. Science Immunology 8, eadg2979 (2023).

9. Von Niessen, A. G. O. et al. Improving mRNA-based therapeutic gene delivery by expression-augmenting 3’ UTRs identified by cellular library screening. Molecular Therapy 27, 824–836 (2019).

10. Xia, Z. et al. Dynamic analyses of alternative polyadenylation from RNA-seq reveal a 3’-UTR landscape across seven tumour types. Nature communications 5, 5274 (2014).

11. Ha, K. C., Blencowe, B. J. & Morris, Q. QAPA: a new method for the systematic analysis of alternative polyadenylation from RNA-seq data. Genome biology 19, 1–18 (2018).

12. Derti, A. et al. A quantitative atlas of polyadenylation in five mammals. Genome research 22, 1173–1183 (2012).

13. Shepard, P. J. et al. Complex and dynamic landscape of RNA polyadenylation revealed by PAS-Seq. Rna 17, 761–772 (2011).

14. Hoque, M. et al. Analysis of alternative cleavage and polyadenylation by 3’ region extraction and deep sequencing. Nature methods 10, 133–139 (2013).

15. Moll, P., Ante, M., Seitz, A. & Reda, T. QuantSeq 3’ mRNA sequencing for RNA quantification 2014.

16. Zheng, G. X. et al. Massively parallel digital transcriptional profiling of single cells. Nature communications 8, 14049 (2017).

17. Macosko, E. Z. et al. Highly parallel genome-wide expression profiling of individual cells using nanoliter droplets. Cell 161, 1202–1214 (2015).

18. Hashimshony, T., Wagner, F., Sher, N. & Yanai, I. CEL-Seq: single-cell RNA-Seq by multiplexed linear amplification. Cell reports 2, 666–673 (2012).

19. Shulman, E. D. & Elkon, R. Cell-type-specific analysis of alternative polyadenylation using single-cell transcriptomics data. Nucleic acids research 47, 10027–10039 (2019).

20. Patrick, R. et al. Sierra: discovery of differential transcript usage from polyA-captured single-cell RNA-seq data. Genome biology 21, 1–27 (2020).

21. Gao, Y., Li, L., Amos, C. I. & Li, W. Analysis of alternative polyadenylation from single-cell RNA-seq using scDaPars reveals cell subpopulations invisible to gene expression. Genome research 31, 1856–1866 (2021).

22. Li, W. V., Zheng, D., Wang, R. & Tian, B. MAAPER: model-based analysis of alternative polyadenylation using 3’ end-linked reads. Genome biology 22, 222 (2021).

23. Wu, X., Liu, T., Ye, C., Ye, W. & Ji, G. scAPAtrap: identification and quantification of alternative polyadenylation sites from single-cell RNA-seq data. Briefings in Bioinformatics 22, bbaa273 (2021).

24. Li, G.-W. et al. SCAPTURE: a deep learning-embedded pipeline that captures polyadenylation information from 3’ tag-based RNA-seq of single cells. Genome biology 22, 221 (2021).

25. Zhou, R. et al. SCAPE: a mixture model revealing single-cell polyadenylation diversity and cellular dynamics during cell differentiation and reprogramming. Nucleic Acids Research 50, e66–e66 (2022).

26. Kang, B. et al. Infernape uncovers cell type–specific and spatially resolved alternative polyadenylation in the brain. Genome Research 33, 1774–1787 (2023).

27. Fu, R. et al. scraps: an end-to-end pipeline for measuring alternative polyadenylation at high resolution using single-cell RNA-seq. bioRxiv, 2022–08 (2022).

28. Frith, M. C. et al. A code for transcription initiation in mammalian genomes. Genome research 18, 1–12 (2008).

29. Bogard, N., Linder, J., Rosenberg, A. B. & Seelig, G. A deep neural network for predicting and engineering alternative polyadenylation. Cell 178, 91–106 (2019).

30. Alipanahi, B., Delong, A., Weirauch, M. T. & Frey, B. J. Predicting the sequence specificities of DNA-and RNA-binding proteins by deep learning. Nature biotechnology 33, 831–838 (2015).

31. Long, Y. et al. Accurate transcriptome-wide identification and quantification of alternative polyadenylation from RNA-seq data with APAIQ. Genome Research 33, 644–657 (2023).

32. Wang, Y. et al. Single-cell transcriptome analysis reveals differential nutrient absorption functions in human intestine. Journal of Experimental Medicine 217, e20191130 (2019).

33. Blazie, S. M. et al. Comparative RNA-Seq analysis reveals pervasive tissue-specific alternative polyadenylation in Caenorhabditis elegans intestine and muscles. BMC biology 13, 1–21 (2015).

34. Wang, X. et al. Long non-coding RNA ZFAS1 promotes colorectal cancer tumorigenesis and development through DDX21-POLR1B regulatory axis. Aging (Albany NY) 12, 22656 (2020).

35. Wang, J. et al. (Pro) renin receptor promotes colorectal cancer progression through inhibiting the NEDD4L-mediated Wnt3 ubiquitination and modulating gut microbiota. Cell Communication and Signaling 21, 2 (2023).

36. Meyer, A. R., Brown, M. E., McGrath, P. S. & Dempsey, P. J. Injury-induced cellular plasticity drives intestinal regeneration. Cellular and Molecular Gastroenterology and Hepatology 13, 843–856 (2022).

37. Habowski, A. N. et al. Transcriptomic and proteomic signatures of stemness and differentiation in the colon crypt. Communications biology 3, 453 (2020).

38. Huang, K. et al. Cell-type-specific alternative polyadenylation promotes oncogenic gene expression in non-small cell lung cancer progression. Molecular Therapy-Nucleic Acids 33, 816–831 (2023).

39. Dinh, H. Q. et al. Integrated single-cell transcriptome analysis reveals heterogeneity of esophageal squamous cell carcinoma microenvironment. Nature communications 12, 7335 (2021).

40. Huang, Y. & Sanguinetti, G. BRIE2: computational identification of splicing phenotypes from single-cell transcriptomic experiments. Genome biology 22, 251 (2021).

41. Chen, H. et al. S100A14: novel modulator of terminal differentiation in esophageal cancer. Molecular Cancer Research 11, 1542–1553 (2013).

42. Kolfschoten, I. G. et al. A genetic screen identifies PITX1 as a suppressor of RAS activity and tumorigenicity. Cell 121, 849–858 (2005).

43. Li, Y. et al. Immune signature profiling identified predictive and prognostic factors for esophageal squamous cell carcinoma. Oncoimmunology 6, e1356147 (2017).

44. Xu, W. W. et al. Targeting VEGFR1-and VEGFR2-expressing non-tumor cells is essential for esophageal cancer therapy. Oncotarget 6, 1790 (2015).

45. Ray, D. et al. A compendium of RNA-binding motifs for decoding gene regulation. Nature 499, 172–177 (2013).

46. Li, X. et al. Comparative transcriptome characterization of esophageal squamous cell carcinoma and adenocarcinoma. Computational and Structural Biotechnology Journal 21, 3841–3853 (2023).

47. Wei, W. et al. Comprehensive characterization of posttranscriptional impairment-related 3’-UTR mutations in 2413 whole genomes of cancer patients. NPJ Genomic Medicine 7, 34 (2022).

48. Gregersen, L. H. et al. SCAF4 and SCAF8, mRNA anti-terminator proteins. Cell 177, 1797–1813 (2019).

49. He, P. et al. The changing mouse embryo transcriptome at whole tissue and single-cell resolution. Nature 583, 760–767 (2020).

50. Dube, D. K. et al. Qualitative and quantitative evaluation of TPM transcripts and proteins in developing striated chicken muscles indicate TPM4*α* is the major sarcomeric cardiac tropomyosin from early embryonic life to adulthood. Cytoskeleton 75, 437–449 (2018).

51. Li, D., Kishta, M. S. & Wang, J. Regulation of pluripotency and reprogramming by RNA binding proteins. Current topics in developmental biology 138, 113–138 (2020).

52. Kobayashi, T. et al. The cyclic gene Hes1 contributes to diverse differentiation responses of embryonic stem cells. Genes & development 23, 1870–1875 (2009).

53. Li, N. et al. CFIm-mediated alternative polyadenylation safeguards the development of mammalian pre-implantation embryos. Stem Cell Reports 18, 81–96 (2023).

54. Wang, J. et al. Comprehensive mapping of alternative polyadenylation site usage and its dynamics at single-cell resolution. Proceedings of the National Academy of Sciences 119, e2113504119 (2022).

55. Fansler, M. M., Mitschka, S. & Mayr, C. Quantifying 3’ UTR length from scRNA-seq data reveals changes independent of gene expression. Nature Communications 15, 4050 (2024).

56. Li, L. et al. An atlas of alternative polyadenylation quantitative trait loci contributing to complex trait and disease heritability. Nature genetics 53, 994–1005 (2021).

57. Ban, M. et al. Expression profiling of cerebrospinal fluid identifies dysregulated antiviral mechanisms in multiple sclerosis. Brain 147, 554–565 (2024).

58. Qin, S. et al. mRNA-based therapeutics: powerful and versatile tools to combat diseases. Signal transduction and targeted therapy 7, 166 (2022).

59. Dobin, A. et al. STAR: ultrafast universal RNA-seq aligner. Bioinformatics 29, 15–21 (2013).

60. Li, H. et al. The sequence alignment/map format and SAMtools. bioinformatics 25, 2078–2079 (2009).

61. Shirley, M. D., Ma, Z., Pedersen, B. S. & Wheelan, S. J. Efficient” pythonic” access to FASTA files using pyfaidx tech. rep. (PeerJ PrePrints, 2015).

62. Avsec, Ž., et al. The Kipoi repository accelerates community exchange and reuse of predictive models for genomics. Nature biotechnology 37, 592–600 (2019).

63. Wolf, F. A., Angerer, P. & Theis, F. J. SCANPY: large-scale single-cell gene expression data analysis. Genome biology 19, 1–5 (2018).

64. Ramírez, F., et al. High-resolution TADs reveal DNA sequences underlying genome organization in flies. Nature communications 9, 189 (2018).

65. Grant, C. E., Bailey, T. L. & Noble, W. S. FIMO: scanning for occurrences of a given motif. Bioinformatics 27, 1017–1018 (2011).

66. Liao, J.-Y. et al. EuRBPDB: a comprehensive resource for annotation, functional and oncological investigation of eukaryotic RNA binding proteins (RBPs). Nucleic acids research 48, D307–D313 (2020).

67. Zhou, Y. et al. Metascape provides a biologist-oriented resource for the analysis of systems-level datasets. Nature communications 10, 1523 (2019).

68. Wu, P. & Wang, W. Distinct 3D contacts and phenotypic consequences of adjacent non-coding loci in the epigenetically quiescent regions. bioRxiv (2023).

