## Supplementary material for "scTail: precise polyadenylation site detection and its alternative usage analysis from reads 1 preserved 3’ scRNA-seq data": This file includes all supplementary figures.

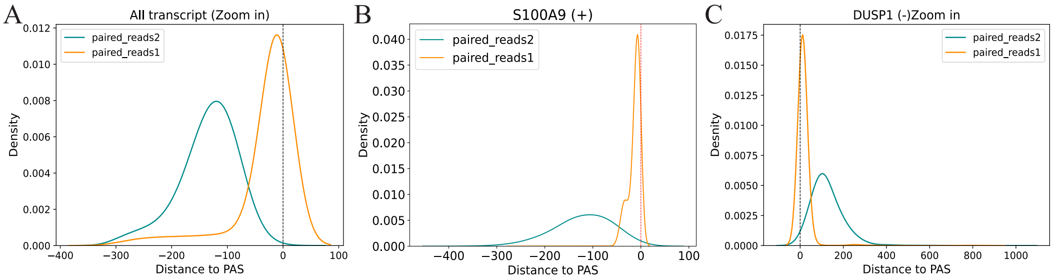


Figure S1: A kernel density estimate (KDE) plot visualize the distribution of the most popular distance between transcript end and reads teminate belong to this transcript for all transcript (left panel), the distribution of distances between reads 1 or reads 2 end and S100A9 gene end (middle panel) and the distribution of distances between reads 1 or reads 2 end and DUSP1 gene end (right panel).

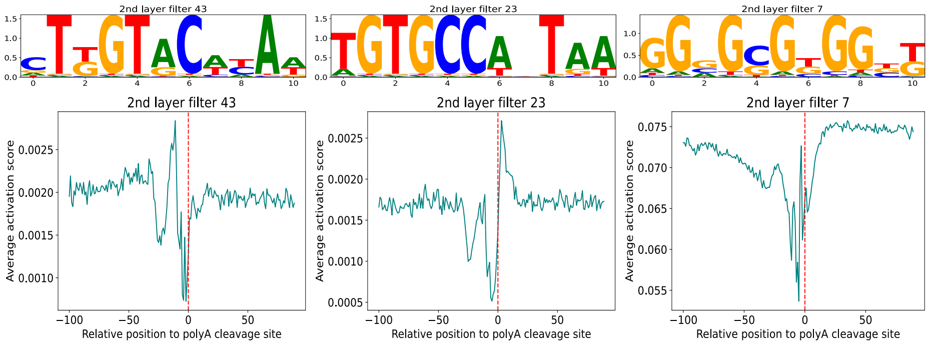


Figure S2: Active motifs of embedded convolutional neural network. The model can identify three characteristic PAS motif, including upstream UGUA motifs, downstream GU-rich and G-rich motifs. Motif activated score was calculated on 200-bp sequences around stringent PASs.

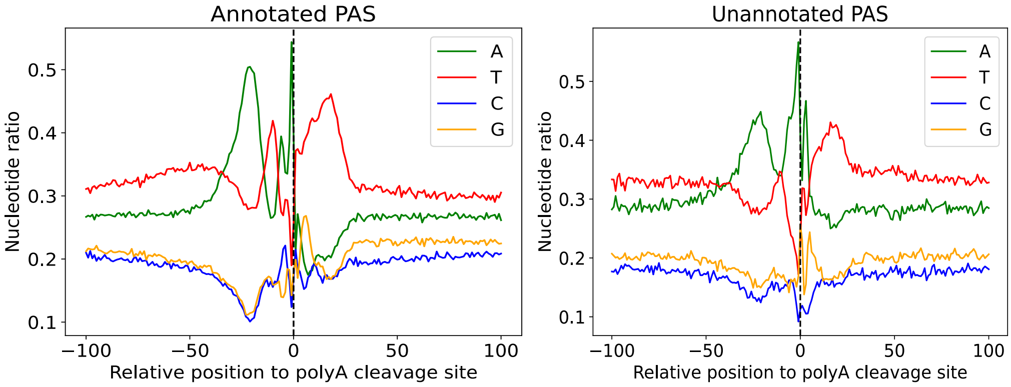


Figure S3: Profile of nucleotide frequencies around annotated (i.e. overlap with GENCODE annoation; left panel) and unannotated (i.e. non-overlap with GENCODE annotation; right panel) PAS identified by scTail. Upstream 100 bp and downstream 100 bp sequences were analyzed.

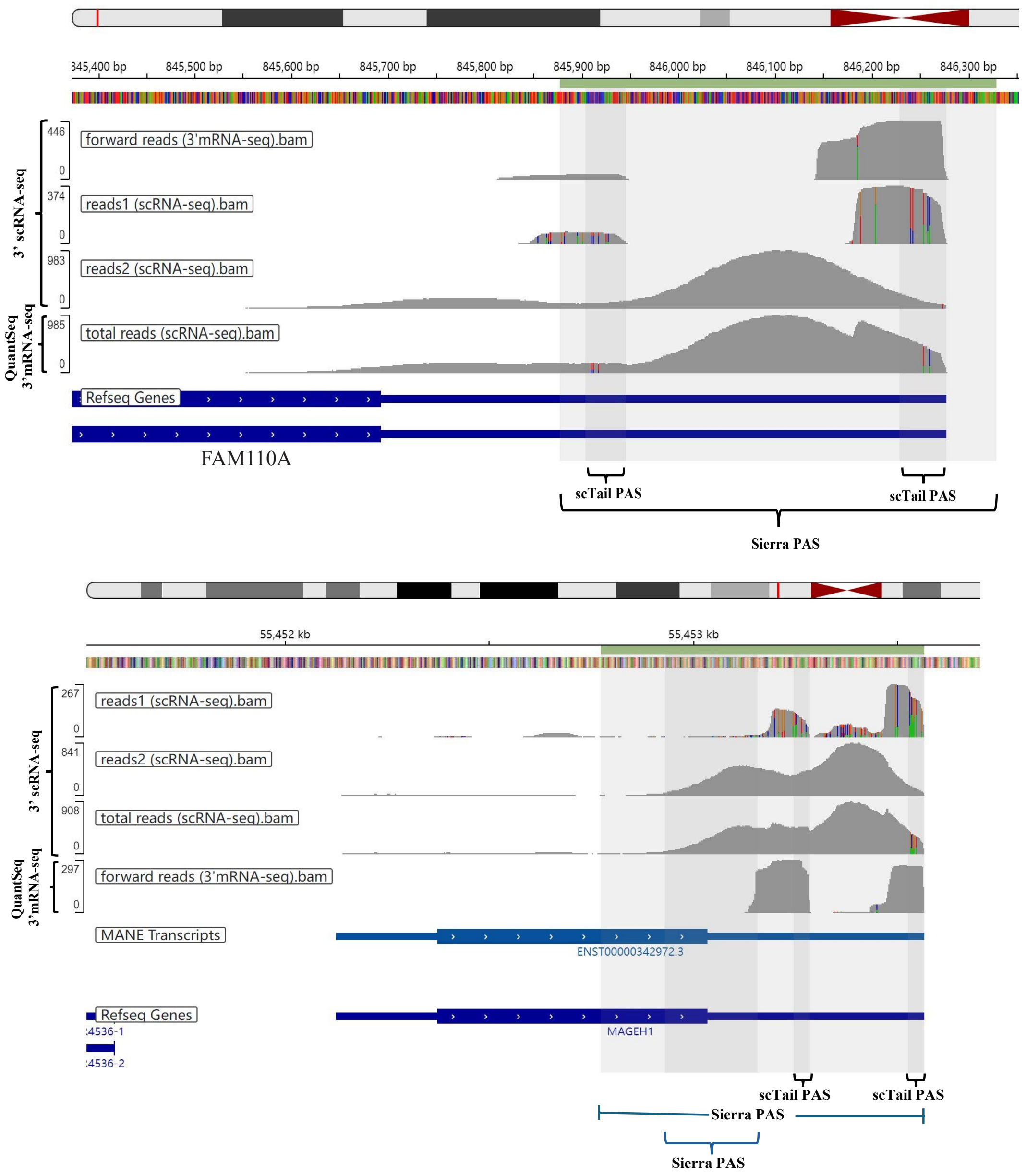


Figure S4: Screenshots from Integrative Genomics Viewer (IGV) shows aligned reads 1, reads 2 and total reads from 3'scRNA-seq (10X Genomics) and forward reads from QuantSeq REV for FAM110A (top panel) and MAGEH1 (bottom panel). Grey regions display PAS detected by scTail and Sierra. The genome position and annotation are shown at the top and bottom, respectively.

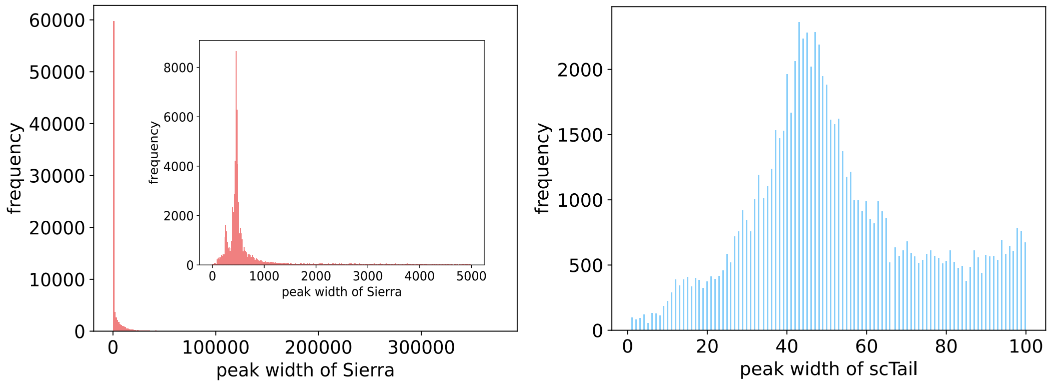


Figure S5: Histogram shows the distribution of peak widths of Sierra (left panel) and scTail (right panel). The small figure within the left panel is the zoom in result of Sierra.

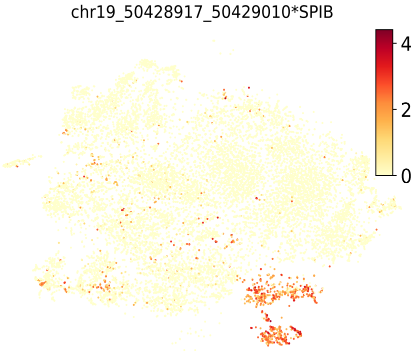


Figure S6: The expression of one PAS marker (i.e. chr19_50428917_50429010*SPIB) of transit-amplifying (TA) cells.

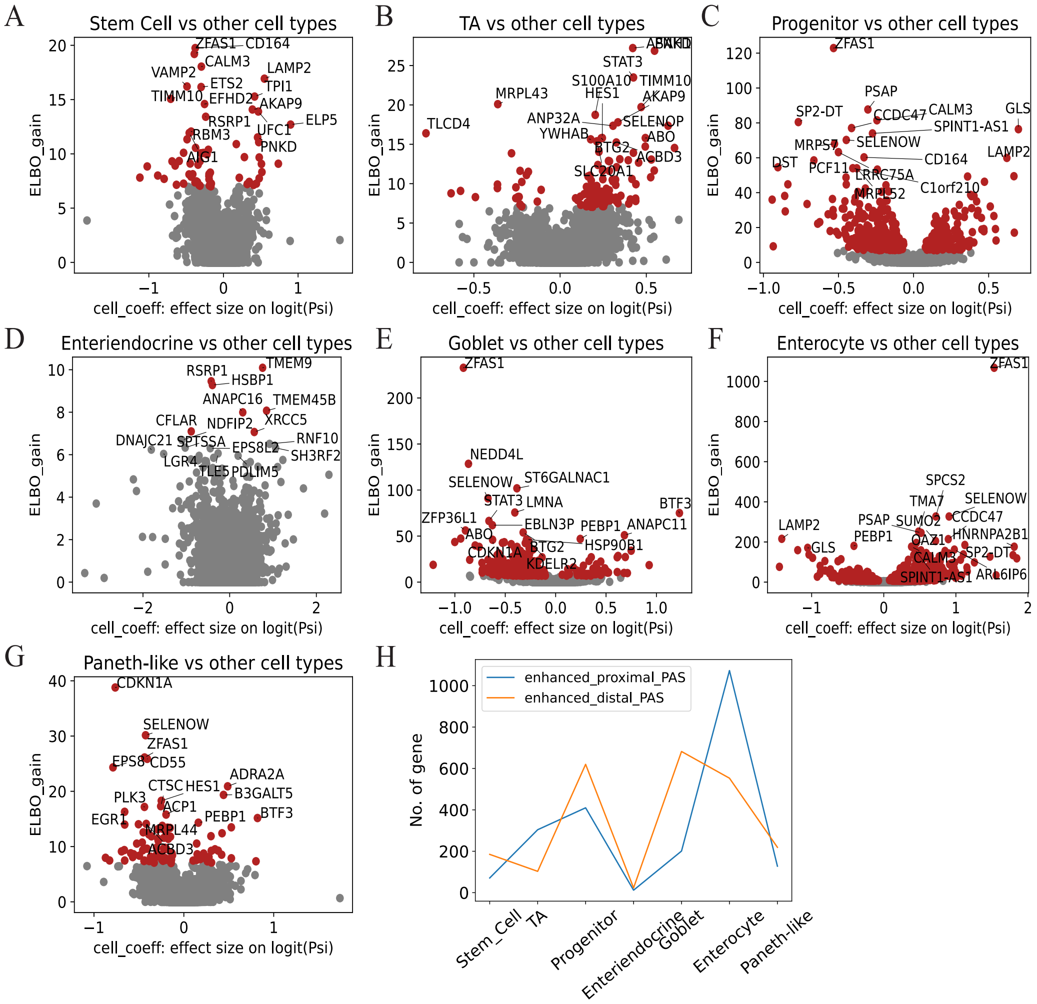


Figure S7: Significant differential genes with alternative polyadenylation site across multiple cell types in human intestinal. Volcano plot between ELBO_gain and effect size on logit (PSI) for detecting differential polyadenylation between stem cell (A), transit-amplifying cell (B), progenitor (C), enteriendocrine (D), goblet (E), enterocyte (F) or paneth-like cell (G) and other cells by BRIE2. Line chart shows the number of genes significantly preferring to use proximal and distal PAS in different cell types (FDR $< 0.01$; H).

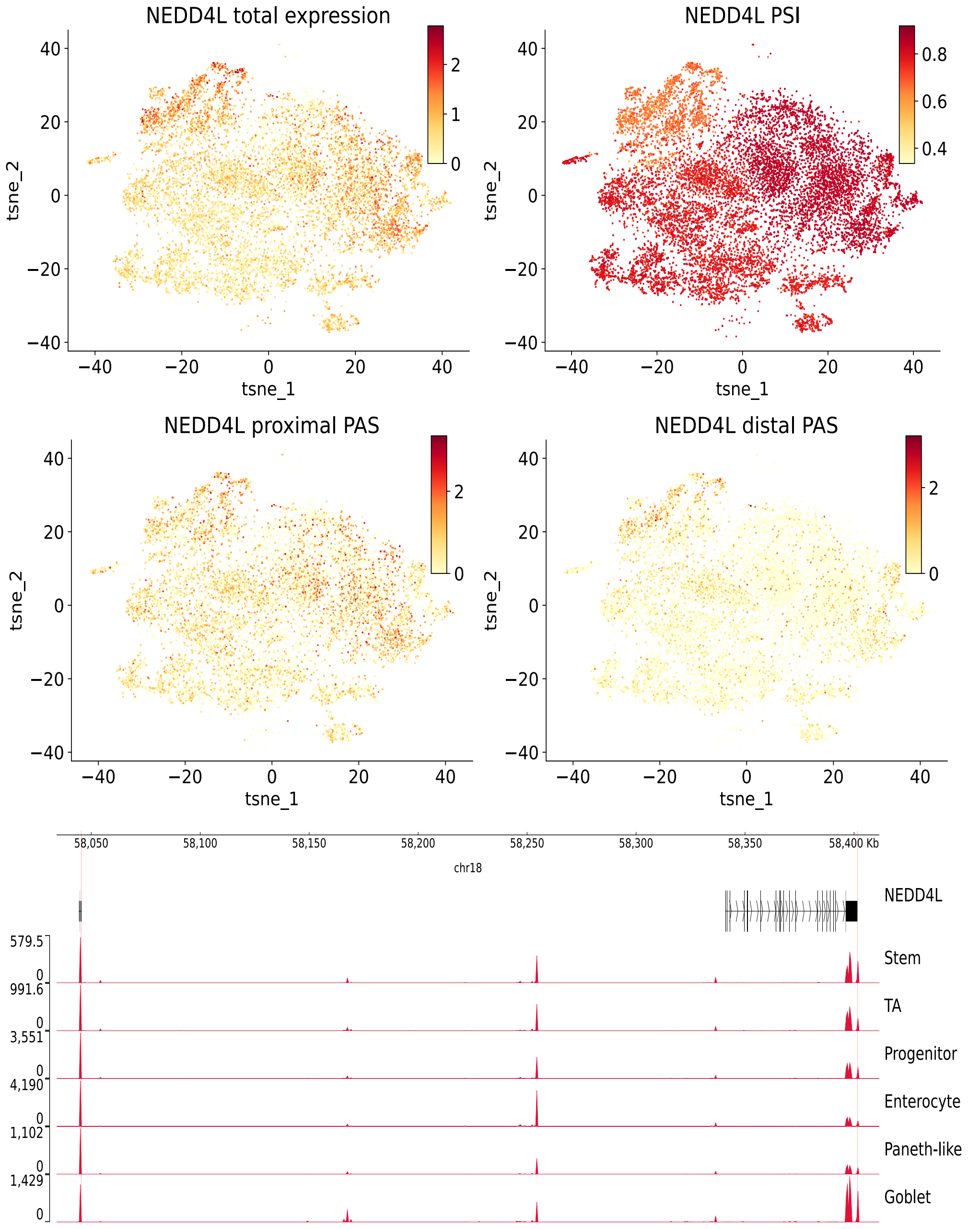


Figure S8: Example gene NEDD4L displays cell-type specific of APA usage. tSNE plot shows total RNA expression, PSI, proximal and distal PAS expression of NEDD4L in all human intestine cells. Genomic track plot illustrates aligned reads coverage of all cell types in human intestinal. Genome annotation of NEDD4L shows two transcripts

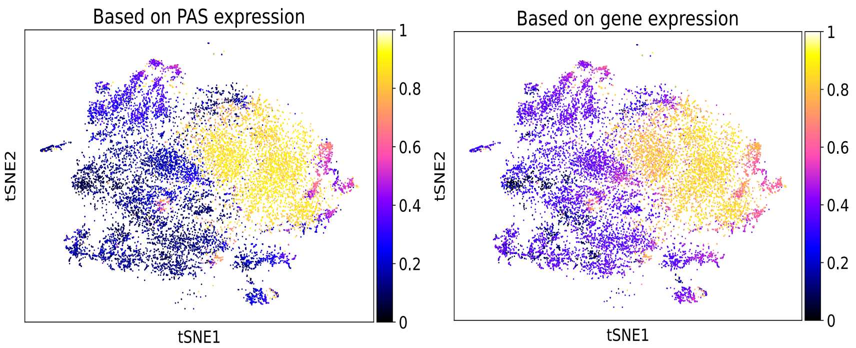


Figure S9: tSNE plots show diffusion pseudotime of individual cells calculated by expression and PAS profile of genes with differential PAS usage across cell types. Stem cell was regarded as the root cell.

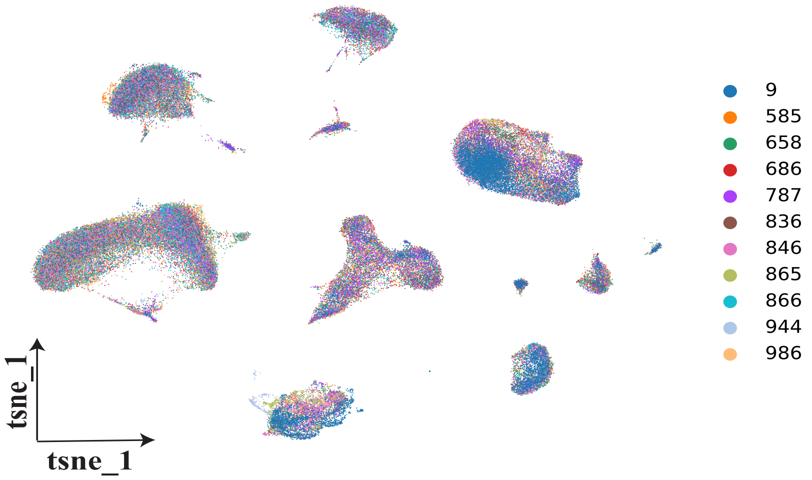


Figure S10: UMAP visualization of all cells (n=89,313) in esophageal squamous cell carcinoma. Each dot represents an individual cell, where colors indicate patient id.

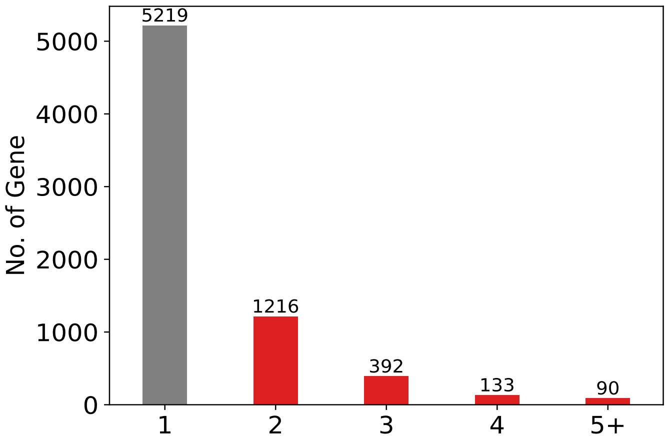


Figure S11: Detected polyadenylation sites across genes: 9,888 PAS annotated to 7,050 genes, 26% of genes possess more than 1 site (red bars) in ESCC dataset.

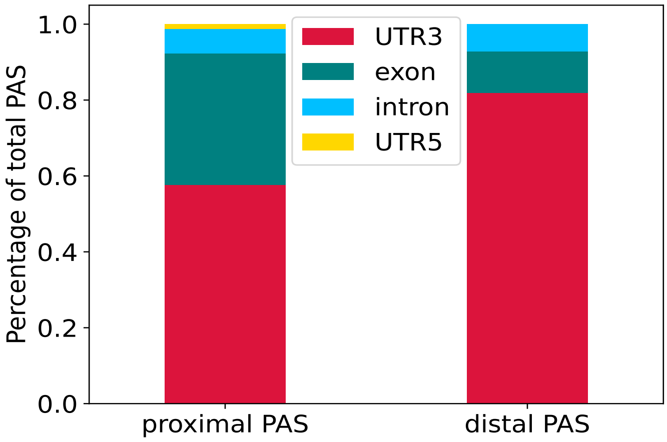


Figure S12: Genomics distribution of proximal and distal site location within the gene.

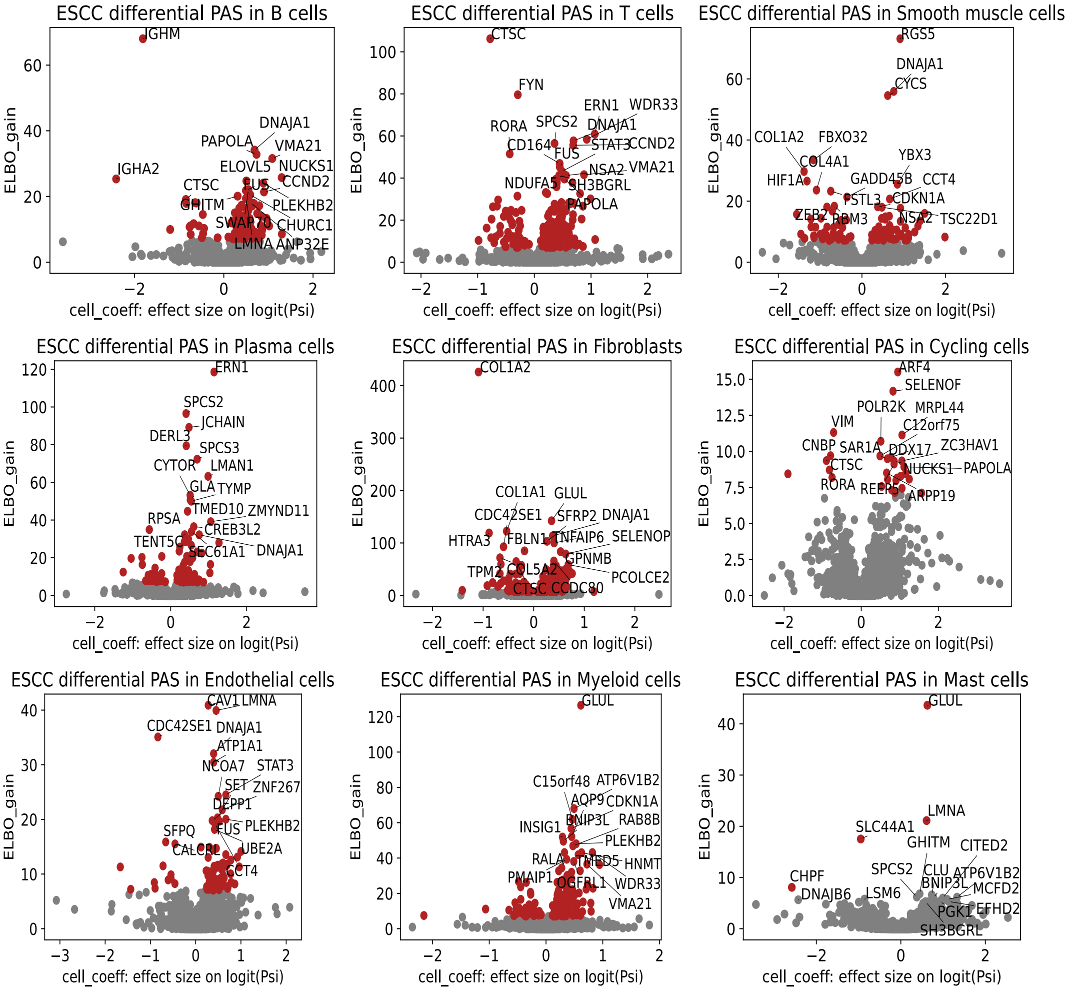


Figure S13: Volcano plot between ELBO_gain and effect size on logit (PSI) for detecting differential polyadenylation site between normal and tumor across B cells, T cells, smooth muscle cells, plasma cells, fibroblasts, cycling cells, endothelial cells, myeloid cells and mast cells in ESCC dataset.

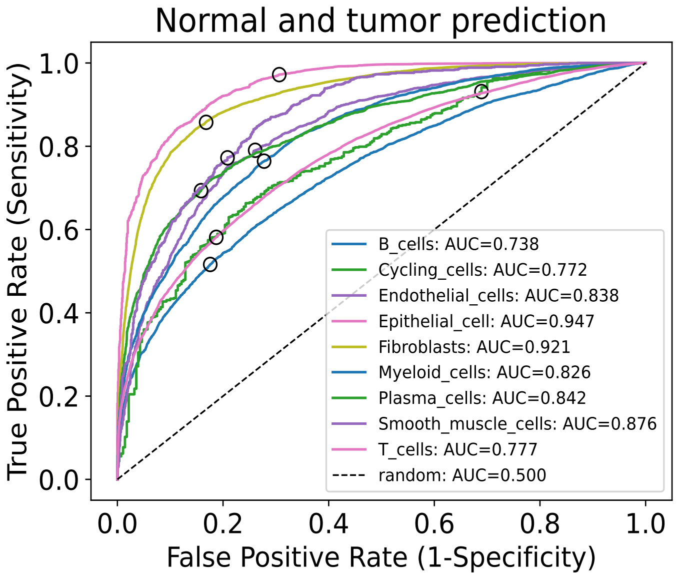


Figure S14: ROC curves for prediction of disease condition from the PSI matrix of significant genes with APA usage by using a random forest in a binary-label classification. Tenfold cross-validation was utilized to evaluate the model.

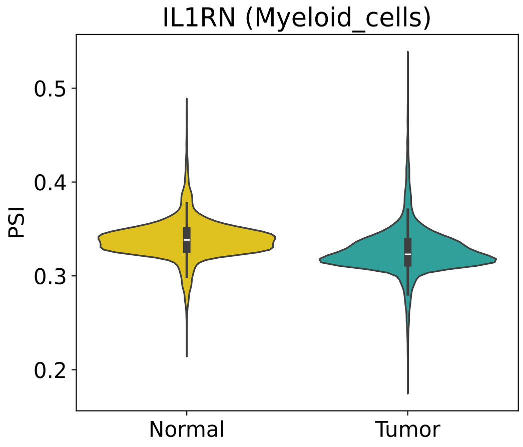


Figure S15: Violin plot shows PSI value of IL1RN in normal and tumor conditions in myeloid cell in ESCC dataset.

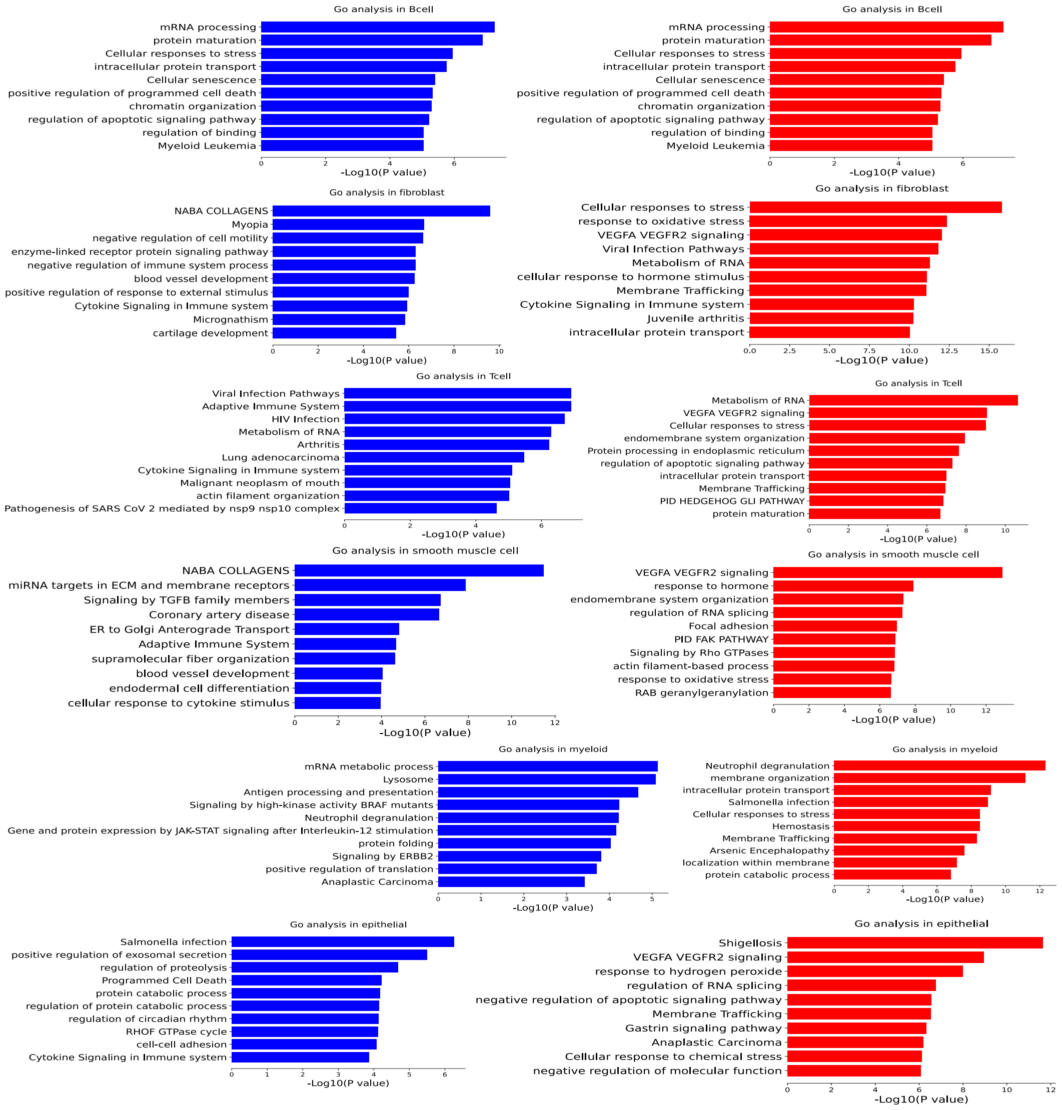


Figure S16: GO terms associated with differential genes preferring to use proximal PAs (blue barplot; left panel) and distal PAS (red barplot; right panel) in tumor condition across B cells, fibroblast, T cells, smooth muscle cell, myeloid and epithelial cell.

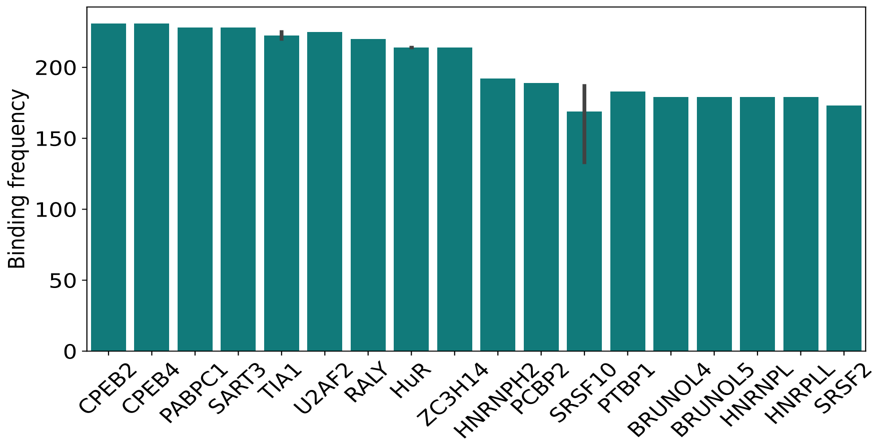


Figure S17: Bar plot shows the binding frequency of RBP with gap sequence detected by FIMO in ESCC dataset.

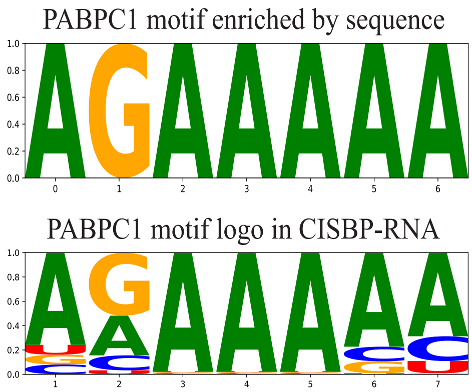


Figure S18: WebLogo of the base frequency of M146_0.6 (i.e. one motif of PABPC1) enriched in the sequences detected by FIMO (top) and displayed in the CISBP-RNA database (bottom).

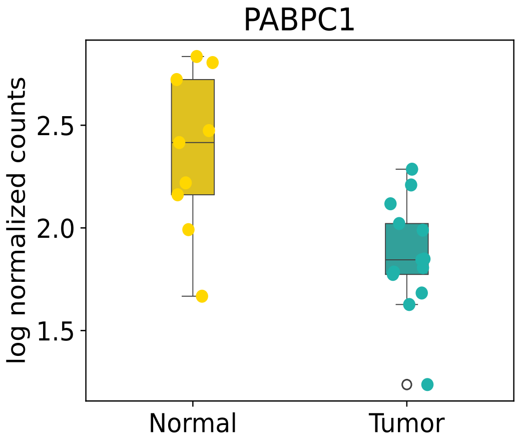


Figure S19: Boxplot displays the log normalized count of PABPC1 in tumor and normal samples. Each dot represent individual sample.

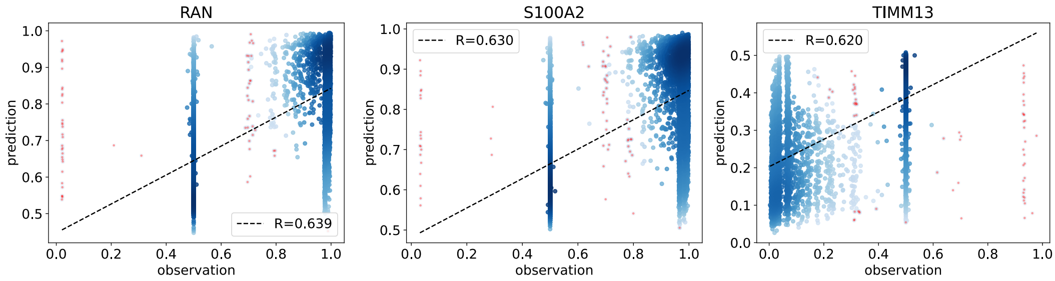


Figure S20: A scatter plot between the predicted with all RBP abundance and measured expression of RAN, S100A2 and TIMM13 in epithelial.

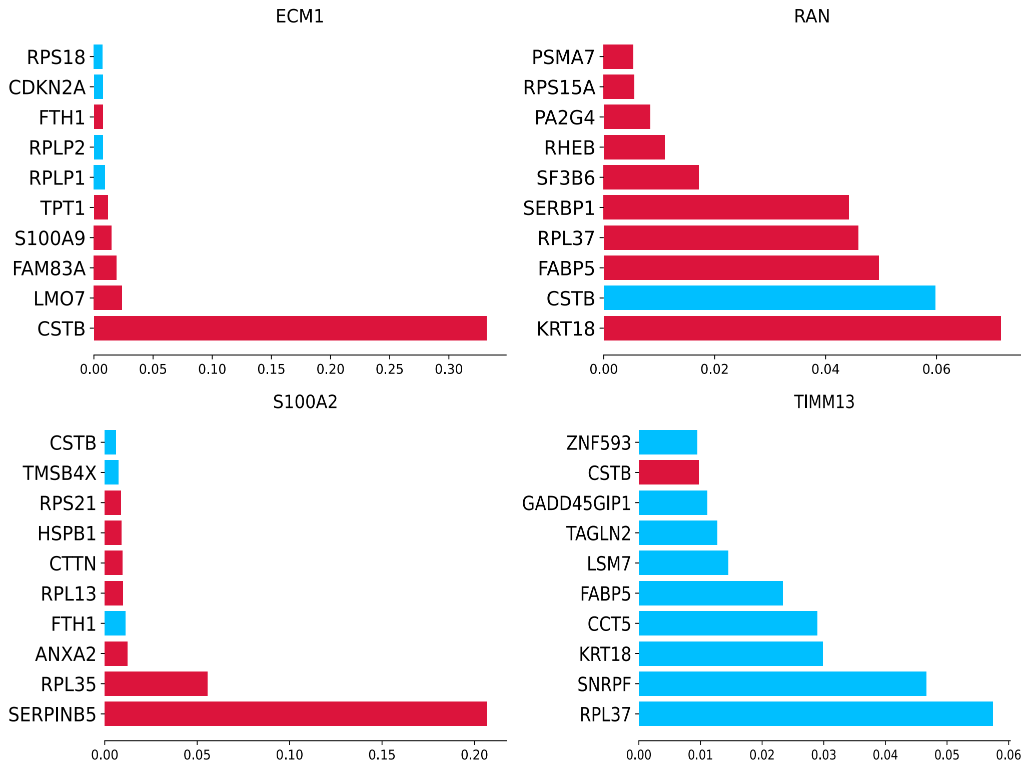


Figure S21: The top 10 important RBPs predicting PSI value of ECM1, RAN, S100A2 and TIMM13. The red and blue bar highlights RBP with positive and negative correlation with PSI, respectively.

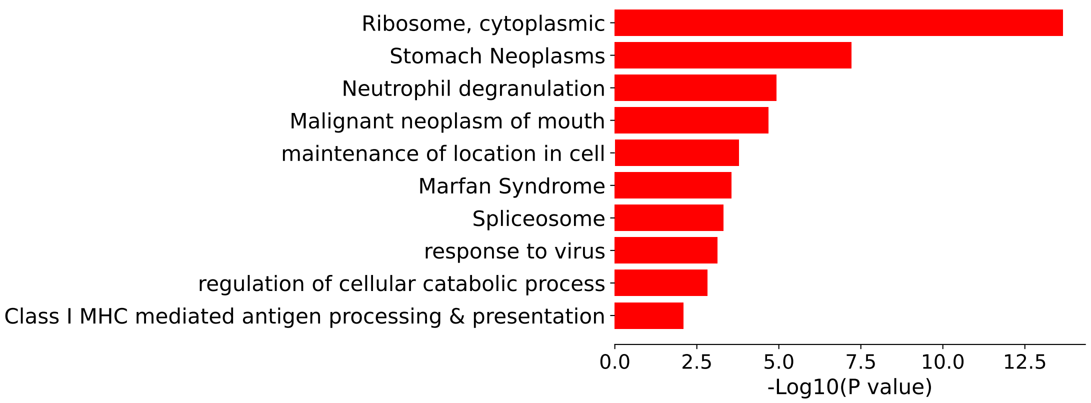


Figure S22: Bar plot shows the enriched terms of important RBP predicting PSI of ECM1, RAN, S100A2 and TIMM13.

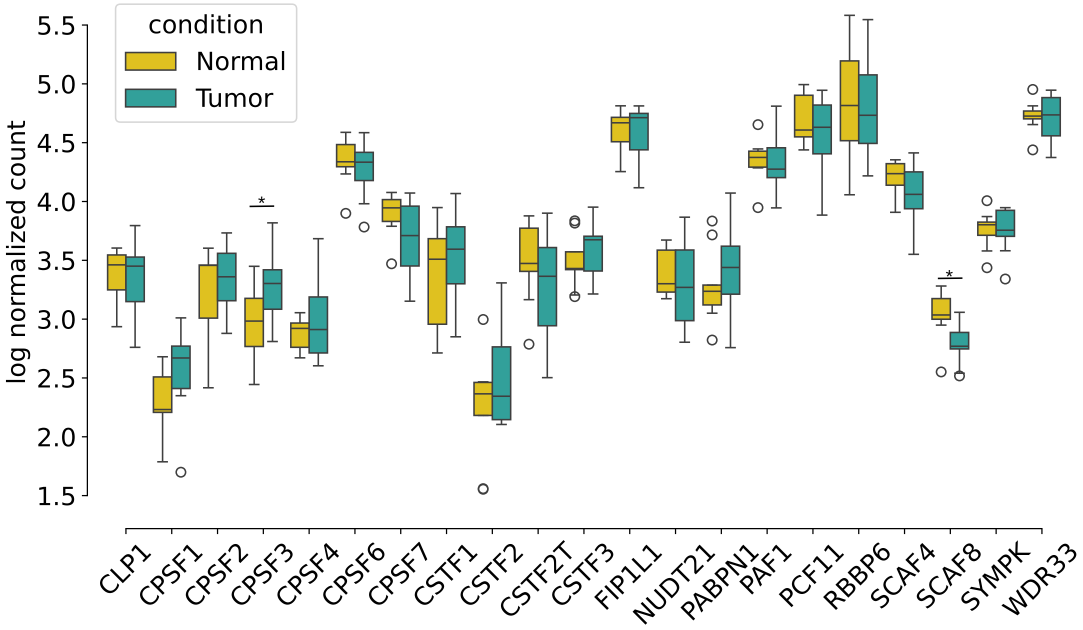


Figure S23: Boxplots show normalized expression of CPA specificity factor (CPSF) in normal and tumor context of esophageal squamous cell carcinoma (ESCC) dataset.

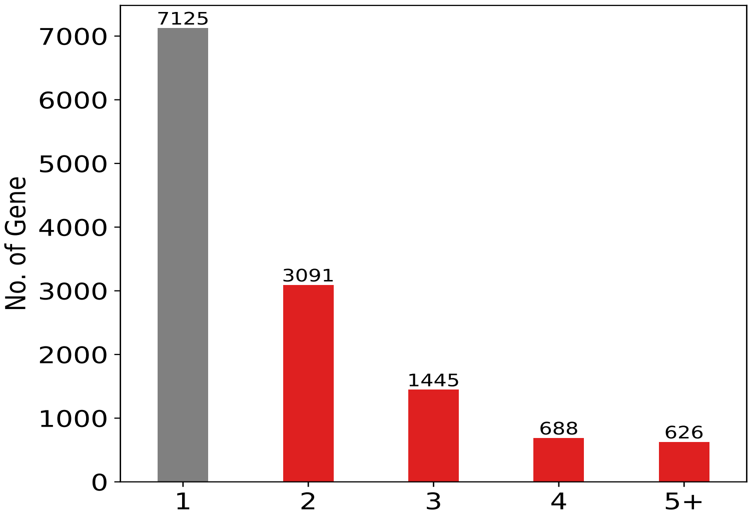


Figure S24: Detected polyadenylation sites across genes: 24,109 PAS annotated to 12,975 genes, 45% of genes possess more than 1 site (red bars) in mouse forelimb development dataset.

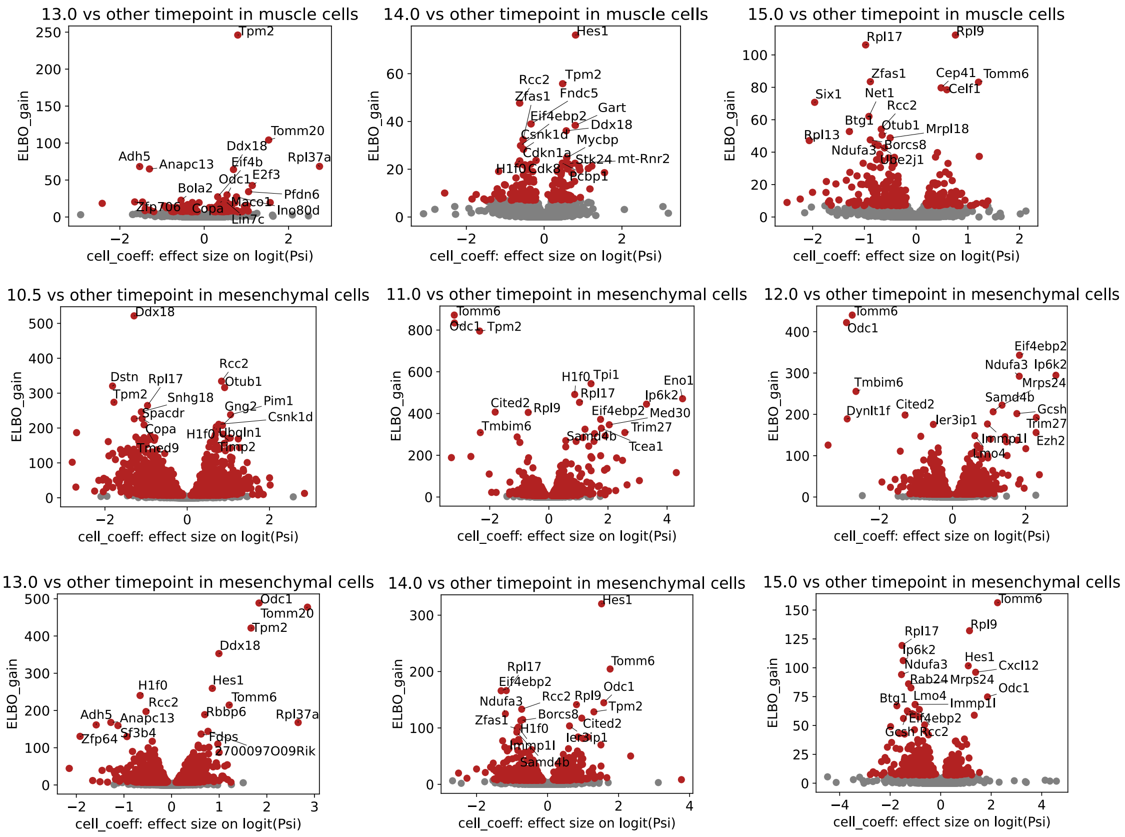


Figure S25: Volcano plots display the correlation between ELBO_gain and effect size on logit (PSI) for identifying differential PAS between E13.0, E14.0 or E15.0 and other time points in muscle cell (top panel), detecting E10.5-, E11.0-, E12.0-, E13.0-, E14.0- and E15.0-specific PAS in mesenchymal cell.

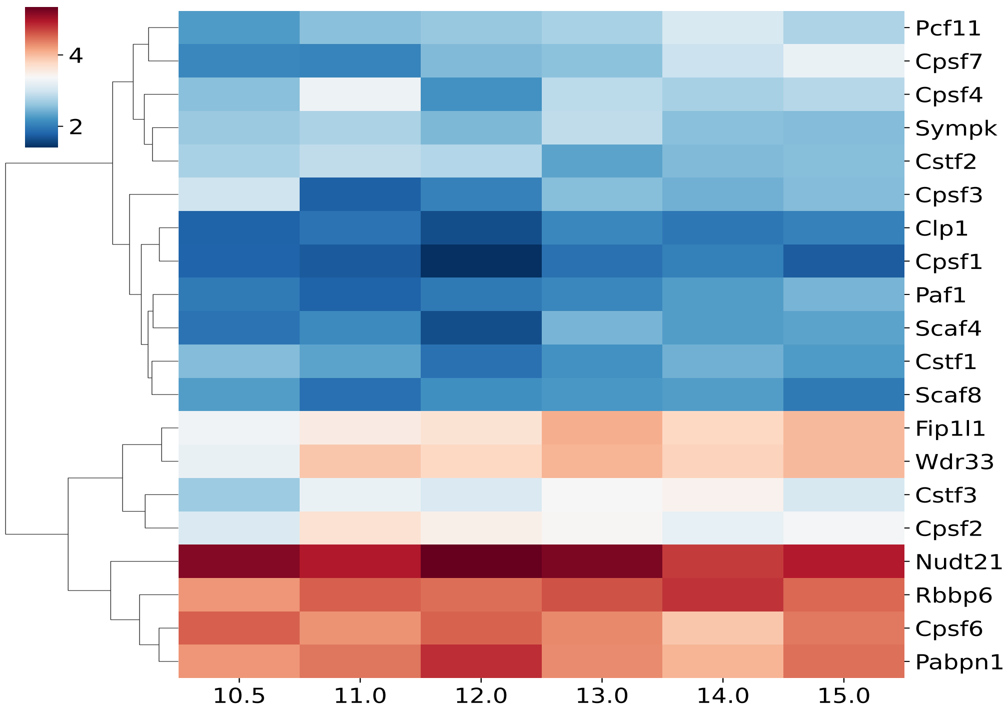


Figure S26: Heatmap shows the normalized expression values of reported CPSF across distinct development stages. Hierarchical cluster the similar expression profiles of CPSFs together.

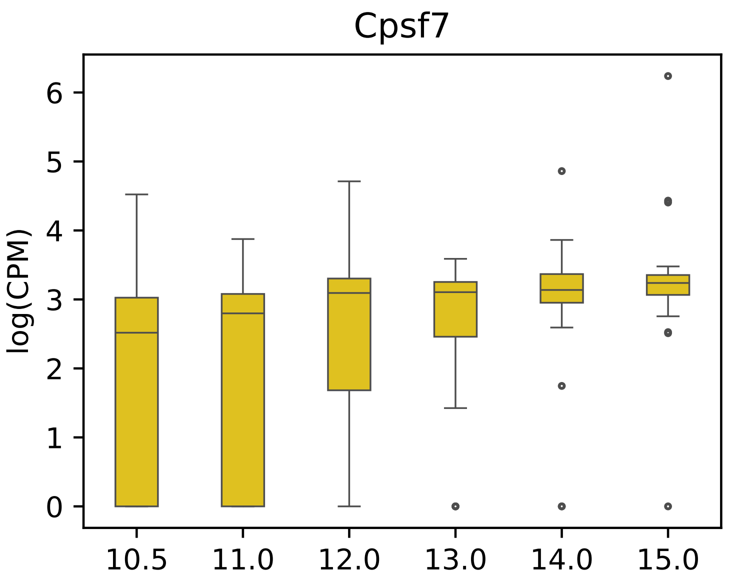


Figure S27: Boxplot shows the expression of Cpsf7 (the only one statistically significant differential gene) across various development stages.
